# TELOMERASE DEPLETION ACCELERATES AGEING OF THE ZEBRAFISH BRAIN

**DOI:** 10.1101/2022.05.24.493215

**Authors:** Raquel R. Martins, Savandara Besse, Pam S. Ellis, Rabia Sevil, Naomi Hartopp, Catherine Purse, Georgia Everett-Brown, Owain Evans, Nadiyah Mughal, Mina HF Wahib, Zerkif Yazigan, Samir Morsli, Ada Jimenez-Gonzales, Andrew Grierson, Heather Mortiboys, Chrissy Hammond, Michael Rera, Catarina M. Henriques

## Abstract

Decreased telomerase expression, telomere shortening, senescence-associated markers and inflammation have all been independently observed in the ageing brain and associated with disease. However, causality between limited telomerase expression and brain senescence and neuro-inflammation in the natural ageing setting is yet to be established. Here, we address these questions using the zebrafish as an ageing model which. Akin to humans, zebrafish display premature ageing and death in the absence of telomerase and where telomere shortening is a driver of cellular senescence.

Our work shows for the first time that telomerase deficiency (*tert^-/-^)* accelerates key hallmarks of ageing identified in the Wild Type (WT) zebrafish brain at transcriptional, cellular, tissue and functional levels. We show that telomerase depletion accelerates ageing-associated transcriptomic changes associated with dysregulation of stress response and immune genes. These are accompanied by accelerated *in situ* accumulation of senescence-associated markers and inflammation in the aged brain. Importantly, *In vivo*, these changes correlate with increased blood-brain barrier permeability and increased anxiety-like behaviour. Of note, the acceleration of senescence-associated markers in the absence of tert occurs not only in the expected proliferative areas but also in non-proliferative ones, where it is unlikely due to telomere-dependent replicative exhaustion, suggesting that non-canonical roles of telomerase may be involved.

Together, our work suggests that telomerase has a protective role in the zebrafish brain against the accumulation of senescence and neuro-inflammation and is required for blood-brain barrier integrity.

**GRAPHICAL ABSTRACT:** Graphical Abstract
Telomerase depletion accelerates markers of ageing in the zebrafish brain, ranging from dysregulated immune and stress ageing transcriptomic hallmarks to *in situ* accumulation of senescence-associated markers and inflammation, dysfunction of the blood-brain barrier and increased anxiety behaviour.

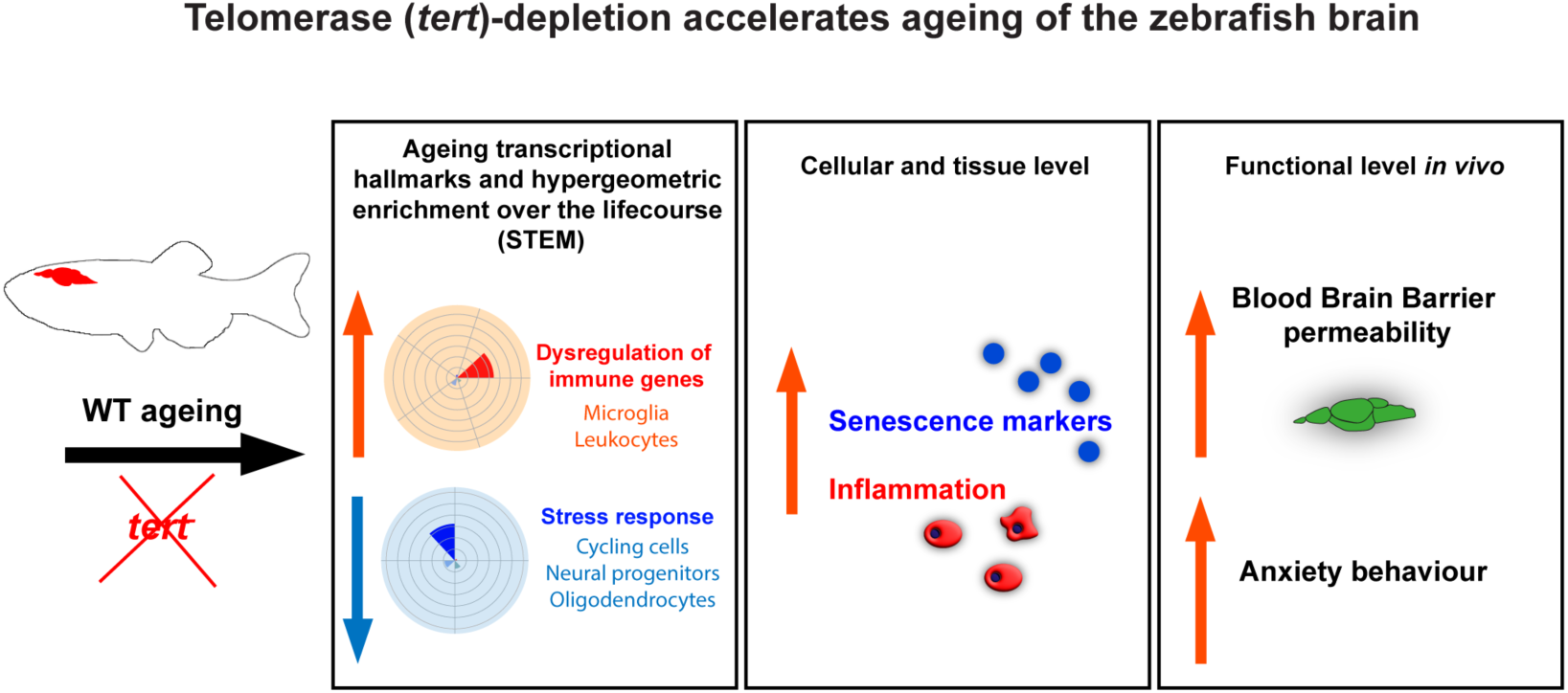

## 1 INTRODUCTION

Brain ageing and related neurodegenerative diseases have dramatic consequences on quality of life and medical care costs. Dementia alone, one of the most prevalent manifestations of neurodegenerative disease affects 23 million people in the UK alone at a cost of around £26 billion a year to the NHS ^1–3^. Understanding the mechanisms underlying brain ageing will target multiple age-associated brain diseases rather than one disease, aiming to ameliorate quality of life in the elderly.

There are well-known key hallmarks of ageing which include telomere attrition, cellular senescence, stem cell exhaustion and chronic inflammation amongst others^4, 5^. Telomeres are (TTAGGG)n DNA repeats that together with a complex of proteins (known as Shelterin) create a “cap-like” structure at the end of linear chromosomes^6^, preventing the ends of linear chromosomes from being recognised as deleterious DNA double strand breaks^7^. However, in humans, due to time- and cell-specific-limited telomerase expression, telomeres shorten with each cell division, eventually leading to proliferative exhaustion and replicative senescence^8, 9^. Even though senescent cells do not proliferate, they are metabolically active, releasing several inflammatory factors, tissue remodelling factors and growth factors, known as Senescence Associated Secretory Phenotype (SASP)^10^. Consequently, accumulation of senescent cells can contribute to impairment of tissue homeostasis, a low level of chronic inflammation and have been linked to several age-associated diseases^11, 12^. Importantly, cellular senescence accumulates with ageing in humans, and this can be thought of as a general stress response^13–15^.

Restricted telomerase expression and consequent telomere dysfunction function are key hallmarks of natural ageing in humans, underpinning multiple age-related diseases^16^. As tissues with high cellular turnover present accelerated telomere erosion^8, 17^, it is reasonable to speculate that telomerase functions are likely to primarily affect highly proliferative tissues. Accordingly, premature accumulation of critically short telomeres has been identified in highly proliferative tissues such as the gut, in tert-deficient animal models, including first generation zebrafish *tert^-/-^* mutants (and late generation TERT KO mice)^17–20^. Nevertheless, because senescence has been shown to “spread”, in a phenomenon described as paracrine senescence, telomere-dependent genome instability and replicative senescence may lead to non-cell autonomous increase in senescence-associated markers and inflammation^21^. This means that the role of telomerase and telomeres is not restricted to highly proliferative tissues. To complicate things further, growing evidence suggests that the Tert component of telomerase also has non-canonical activities, independent of its action at telomeres^22–25^. In the nucleus, these non-canonical functions include transcriptional regulation of genes involved in inflammation, including nuclear factor kappa β (NFkβ) and tumour necrosis factor alpha (TNFα)^26–28^ as well as genes involved in cell proliferation^29, 30^ and cell survival^31, 32^. Telomerase can also translocate to the mitochondria, where it has been shown to play a protective role against DNA damage and oxidative stress^33, 34^. In the brain, considered a predominantly post-mitotic tissue, telomerase has been shown to have a protective role against excitotoxicity^35^, oxidative stress^36^ and neuronal death^37^ all involved in neurodegenerative diseases. Studies in late-generation telomerase-deficient mice have suggested that limited telomerase expression is associated with premature accumulation of senescence-associated markers in different cell populations in the CNS, including Purkinje neurons, cortical neurons and microglia^38–41^. However, it is still unclear whether this is due to telomere length maintenance or non-canonical functions of telomerase^29, 30^.

There are many different stimuli that can cause cellular senescence, including epigenetic alterations, oxidative stress, mitochondrial dysfunction, inactivation of tumour suppressor genes, mechanical or shear stress, pathogens, and activation of oncogenes^42, 43^. What causal role telomerase and telomere dysfunction plays in CNS senescence in humans is, however, not clear, even if implicated^44^. Nevertheless, telomere shortening has been associated with neurodegenerative diseases, including Parkinson’s and AD^45^, including via glia activation and pathogenesis^46^. Of note, a recent study analysing human brain biobank data from over 30,000 patients identified a positive correlation between shorter leukocyte telomere length (used as a proxy for the patient’s general telomere length) and neurodegenerative phenotypes^47^. Additionally, cellular senescence-associated markers have been reported in human neuroinflammation and neurodegenerative diseases. Increased levels of DNA damage checkpoints such as p53, p21 and p16, as well as SASP factors have been observed in the spinal cord and brain of patients with amyotrophic lateral sclerosis (ALS)^38^ and Alzheimer’s disease (AD)^48^, respectively. Higher mRNA levels of p16INK4a in the substantia nigra pars compacta (SNpc) were also observed in post-mortem tissue of patients with Parkinson’s disease (PD) compared to healthy controls^49^. Moreover, senescence, neuroinflammation and oligomeric tau have been associated with AD pathology, recently reviewed here^50^. Hence, growing evidence suggest that, despite being a low proliferative tissue, the CNS accumulates senescence or at least senescence-associated markers, and that this may be associated with brain diseases of ageing. However, causality between limited telomerase expression and increased senescence and neuro-inflammation in the natural ageing setting is yet to be established. Nevertheless, reducing putative senescent cells leads to improvement of cognitive decline in aged mice, suggesting an active role for senescence in this phenotype^51^. It remains to be determined which cell populations undergo senescence in natural ageing, in the CNS, and what type of senescence this is, i.e., replicative (telomere-dependent) or stress-induced senescence. Understanding whether and how telomerase restriction may contribute to ageing-associated brain degenerative phenotypes would help understand the fundamental mechanisms of brain functional decline and potentially contribute to the uncovering of new therapeutic targets to delay or prevent disease.

The role of telomerase restriction and its consequences in natural ageing may benefit from additional and complementary models, beyond the mouse, where most lab-strains have significant longer telomeres than humans and show little evidence of replicative senescence^52, 53^. The zebrafish may offer such a model since, like in humans, telomerase depletion leads to significant decrease in health and lifespan^18–20, 54^. As it was shown that the zebrafish brain presents low levels of Tert in the adult brain and that these decrease further with ageing^55, 56^, we hypothesised that limited expression of Tert in the aged zebrafish brain contributes to brain degeneration with ageing. To our knowledge, this manuscript represents the first broad characterisation of zebrafish brain ageing in the presence and absence of telomerase (Tert).

To test our hypothesis, we performed RNA sequencing in whole brains of zebrafish throughout their lifecourse in both the presence and absence of telomerase (tert). Specifically, we compared WT and *tert^-/-^* siblings derived from tert^+/-^ in-crosses at the age ranges of 2-6, 9-16 and 18-24 for WT and *tert^-/-^* and up to 30-36 months in WT only, since the *tert^-/-^* do not live past 18-24 months. *tert^-/-^* zebrafish, extensively characterised elsewhere ^18, 19, 57^, display no telomerase activity and have significantly shorter telomeres from birth, consequently ageing and dying prematurely. Ageing is usually described as a time-dependent change in tissue homeostasis, that increases the probability of disease and death^58^. However, whether the genes and pathways driving or accompanying these time-dependent changes are also consistently changing in a time-specific manner, remains unresolved ^59^. We therefore decided to combine a time-series analysis (STEM), which allowed the identification of genes and pathways that are consistently up or down-regulated over-time, with the more traditional differential gene expression (DEGs) analysis between young and old animals. This combined strategy allowed the identification of genes that change in a monotonic, time-dependent manner (STEM), versus genes that change at specific stages of life (DEGs). Both transcriptomic analyses strategies, combined with enrichment strategies highlighted increased immune response and decreased proliferation as the main signatures of ageing in both WT and *tert^-/-^* zebrafish brains. We then asked which specific WT ageing-associated gene changes (STEM and standard DEG comparisons between time points) were accelerated, i.e., occurred at an earlier age, in the absence of telomerase (tert). The more refined STEM-based analyses identified 6 specific gene changes accelerated from the age of 9-16 months in *tert^-/-^* and 11 from the age of 18-24, contrasting to 30-36 months in WT. Broader DEG comparisons-based analyses were used to identify DEGs of “old age” (WT young *VS* old) and then compared with *tert^-/-^* at the different ages to determine how telomerase depletion may accelerate transcriptomic changes of “old age”. This approach identified 4 DEGs accelerated from the early age of 2-6 months in *tert^-/-^*; 639 from the age of 9-16 and 109 from the age of 18-24. These transcriptomic changes were accompanied by an *in situ* acceleration of senescence-associated markers and inflammation. *In vivo*, telomerase depletion accelerated ageing-associated increased in blood-brain barrier permeability and increased anxiety-like behaviour. Together, our work suggests that telomerase has a protective role in the zebrafish brain against the accumulation of senescence and neuro-inflammation and is required for BBB integrity.

## 2 RESULTS

### 2.1 Telomerase (tert) depletion accelerates transcriptomic changes of ageing in the zebrafish brain

Tissue-specific transcriptomic analysis over the life course can offer important insights into the downstream molecular mechanisms underpinning age-associated pathologies. Significant research is being dedicated to these approaches in different animal models, including in mice^60–62^. To identify telomerase-dependent and -independent transcriptional signatures of ageing in the zebrafish brain, we performed RNA-Sequencing of whole brain tissue, throughout the lifespan of WT and telomerase-deficient (*tert^-/-^*) fish (**Fig. 1** and supplementary source data files). While *tert^-/-^* fish have a lifespan of c. 18-24 months, WT fish typically die between c. 36-42 months of age ^20^. The data shown here include 4 age-groups of WT (2-6, 9-16, 18-24 and 30-36 months), corresponding to young, adult, median lifespan and old, respectively. As the *tert^-/-^* fish have a shorter lifespan compared with their WT siblings, the data include 3 age-groups of telomerase-deficient fish (2-6, 9-16, and 18-24 months), which correspond to young, medium lifespan and old (see Materials & Methods for further explanation of genotypes and ages chosen) (**Fig 1A**). The reads were aligned to the latest zebrafish genome build GRCz11^63^ and resulted in uniquely mapped read percentages ranging from 92.1% to 94.4%, which is considered good quality^63^. All samples had at least 10 million uniquely mapped reads. To analyse the overall impact of the genotype and age on transcriptomic regulation, we performed a Principal Component Analysis (PCA) and observed that the samples cluster mainly *per* age (**Supp Fig 1A**).

**Figure 1.**
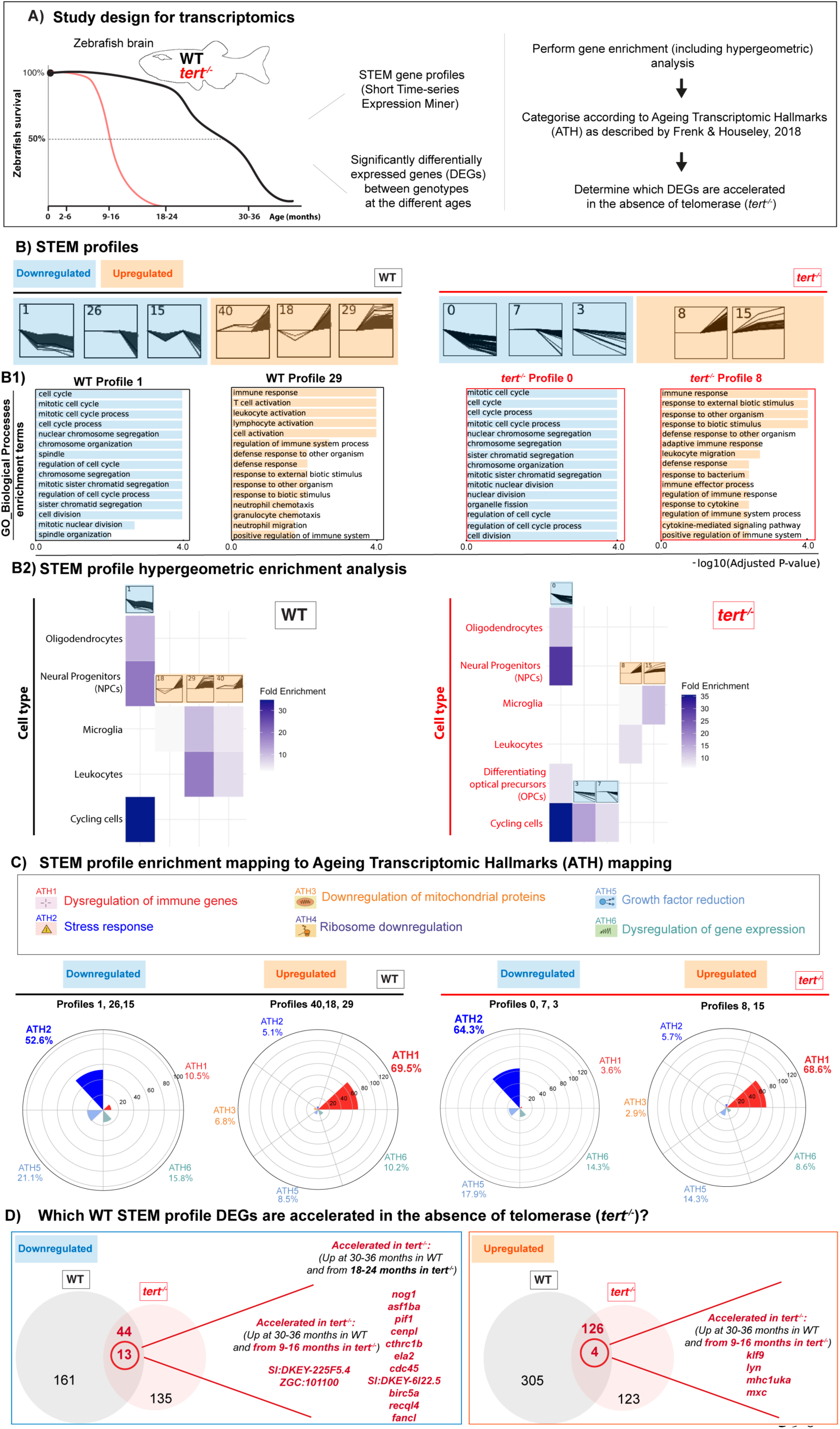
Transcriptomic signatures of ageing in the presence and absence of telomerase in the zebrafish brain. (A) Schematic figure of the study design for transcriptomics: RNA from whole brain tissue from young (ca. 2-6 months), young adult (ca. 9-16 months), middle aged (ca. 18-24 months) and old (,30-36 months) WT fish, and young (ca. 2-6 months), middle aged (ca. 9-16 months) and old (ca. 18-24 months) *tert^-/-^* fish was used for RNA sequencing. N=3 per group. Transcriptomic alterations of ageing were analysed using two complementary approaches: STEM profiles and traditional DEG analysis between genotypes at different ages. Enrichment analysis and hypergeometric analysis were performed to identify the predominant pathways altered with ageing in the presence and absence of telomerase, and the DEGs were mapped to the ageing transcriptomic hallmarks (ATH). (B) Plots of the profiles identified by the STEM analysis (profiles containing down-regulated genes in blue and profiles containing up-regulated genes in orange) in both WT and *tert^-/-^* zebrafish brains, and (B1) top enrichment pathways (GOBP) identified in representative profiles. (B2) STEM profile hypergeometric enrichment analysis in WT and *tert^-/-^* brains identified which cell populations are mainly affected with ageing in the presence and absence of telomerase. (C) The genes identified by STEM analysis were then mapped to ATH. Plots show the proportion of genes mapped to each ATH, in both WT and *tert^-/-^* fish. (D) Venn diagrams highlight the genes identified in the STEM profiles of WT fish that are accelerated in the absence of telomerase (i.e. in *tert^-/-^* fish).

We performed a time-series analysis (STEM) (**Fig 1** and supplementary source (SS) data), which allowed the identification of genes and pathways that are consistently up or down-regulated over-time. In parallel, we performed a broader, more traditional differential gene expression (DEGs) analysis at the different time-points, in WT and *tert^-/-^* (**Supp Fig 1** and SS data). For the time-series analysis, we grouped the gene expression data into temporal expression profiles using Short Time-series Expression Miner (STEM) software^64^. To determine whether the temporal profiles were associated with specific biological processes and pathways, pathway over-representation analysis (ORA) were performed for the genes assigned to the significant STEM profiles. In the WT brain, we identified 6 temporal profiles, 3 of them including overall up-regulated genes (profiles 40, 18 and 29), and 3 including overall down-regulated genes (profiles 1, 26 and 15). In the *tert^-/-^* brain, time-series analysis revealed 5 different profiles. Profiles 8 and 15 containing overall up-regulated genes and profiles 0, 7 and 3 containing overall down-regulated genes (**Fig. 1B** and SS data). We then performed two types of enrichment analyses strategies: Gene Ontology Biological Pathway (GOBP) and hypergeometric. GOBP analysis of STEM profiles highlighted up-regulation of immune response and downregulation of cell-cycle and related pathways with ageing in both WT and *tert^-/-^* zebrafish brains (**Fig 1B1**). The hypergeometric enrichment analysis allowed us to further investigate the cellular context of age-associated transcriptional changes. To do this, we mapped the gene sets from significant STEM profiles in WT and *tert^-/-^* mutant brains onto a published zebrafish brain single-cell dataset^65^. This dataset was chosen because it provides high-quality profiles (>16,000 cells at ∼8,000 UMIs per cell), detailed annotations of unsorted cells capturing the full cell repertoire, and accessible raw and processed data. In WT brains, the major downregulated trajectory (STEM profile 1) was most strongly enriched for genes expressed in cycling cells, with lesser enrichment in neural progenitor cells (NPCs), and oligodendrocytes, indicating a decline in proliferative and myelinating populations with age. Similarly, in *tert^-/-^* mutants, multiple downregulated profiles (0, 3, and 7) showed enrichment for cycling cells, NPCs, oligodendrocytes, and additionally in differentiating oligodendrocyte progenitor cells (OPCs). In contrast, upward trajectories in both WTs (profiles 18, 29, and 40) and *tert^-/-^* mutants (profiles 8 and 15), were enriched for leukocytes and microglia, consistent with increased immune activity during ageing (**Fig 1B2**). These enrichment analyses were further supported when we contextualise pathway enrichments within the hallmarks of ageing, using a previously described strategy^66^ (see additional details in Materials and Methods). Here, we mapped known GOBP terms to experimentally identified ageing transcriptomic hallmarks (ATH)^67^. ATH mapping identified dysregulation of immune genes (ATH1) in up-regulated STEM profiles and stress response (ATH2) in down-regulated STEM profiles in both WT and *tert^-/-^* zebrafish (**Fig 1C** and SS data). The broader analysis of “DEGs of old age” (WT young VS old) showed a GOBP enrichment for cell cycle, division, chromatin-organisation and chromosome segregation and downregulation of DNA transcription as the top 5 enriched pathways. Notably, when mapped to ATH, this broader analysis identified dysregulation of gene expression, stress response and dysregulation of immune genes as the top 3 hallmarks of old age^67^ (**Supp Fig 1** and SS data), supporting the STEM analysis. We further validated selected DEGs of old age by RT-QPCR (*ctss2.1* and *parp3*) and/or *in situ* hybridisation (*cdkn2a/b “p16-like* and *cdkn1a “p21-like”*) (**Fig 2B3** and **Supp Fig 4**).

**Figure 2.**
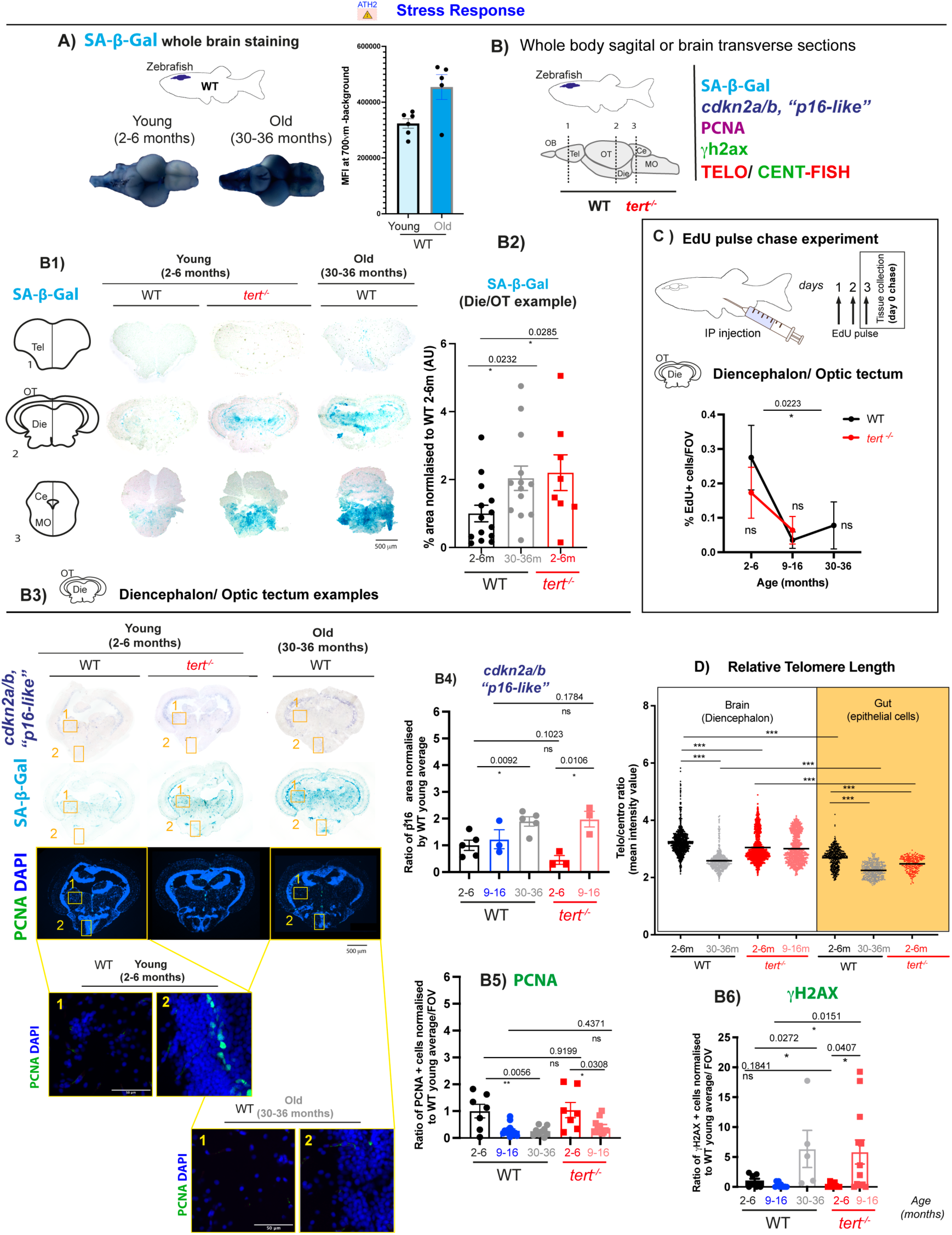
Telomerase (tert) depletion accelerates ageing-associated stress response in the zebrafish brain. Brain tissue collected from ca. 2-6 months, ca. 9-16 months and >30 months) WT fish, and ca. 2-6 months and ca. 9-16 months *tert^-/-^* fish were used to assess stress response by SA-β-Gal staining, *cdkn2a/b* (“*p16-like*”) RNA expression, proliferation (PCNA and EDU staining), telomere length and DNA damage. (A) Whole brain SA-β- Gal staining in young and old WT fish and respective quantification show an increased expression of SA-β-Gal in old WT brains compared to young ones. N=5-6 per group. (B) Schematic figure showing the regions of the brain assessed for senescence-associated markers. (B1) Representative images of SA-β-Gal staining in coronal cryosections of WT and *tert^-/-^* zebrafish show that there is an increased expression of SA-β-Gal in old WT zebrafish (>30 months) and that a similar pattern is observed in *tert^-/-^* at the young age of 2-6 months, particularly in the diencephalon and optic tectum for which (B2) quantifications are shown. N=8-14 per group. (B3) Representative images from *in-situ* hybridisation in coronal cryosections, show an increased expression of *cdkn2a/b* (“*p16-like*”) in WT zebrafish >30 months optic tectum and diencephalon, in areas where SA-β-Gal expression is observed. This increase in *p16* expression is already significant at 9-16 months of age in *tert^-/-^*. Quantifications shown in B4. N=3-5 per group. (B3) Representative images of PCNA and γH2AX staining in adjacent sections from zebrafish brain, in coronal cryosections. (B5) Proliferation assessed by PCNA staining show a decrease in proliferation with ageing in both WT and *tert^-/-^* fish, and this is not accelerated in the absence of telomerase. N=3-7 per group. (B6) DNA damage response assessed by γH2AX staining show increased DNA damage in WT fish >30 months, this increase is anticipated in the *tert^-/-^* fish at 9-16 months but not at 2-6 months of age. N=3-7 per group. (C) Similar to what was observed by PCNA staining, proliferation assessment by EDU staining after a 3-day pulse experiment show a decrease in proliferation with ageing in both WT and *tert^-/-^* brains, but this is not anticipated in the absence of telomerase. N=3-7 per group. (D) Relative telomere length, assessed by telo/cent-FISH, in longitudinal paraffin sections of zebrafish brain (white) and gut (yellow), show that telomere length decreases with ageing in WT fish (>30 months) and that this is anticipated in young *tert^-/-^* fish (ca. 2-6 months). N=4-5 per group. (A, B2, B4, B5, B6) Each dot represents one animal. (D) Each dot represents one cell. Error bars: SEM. * p<0.05; ** p<0.01; *** p<0.001. Abbreviations: OB, olfactive bulb; Tel, telencephalon; Die, diencephalon; OT, optic tectum; Ce, cerebellum; MO, medulla oblongata.

Finally, we asked which specific WT ageing-associated gene changes from the STEM and standard DEG comparisons between time points) were accelerated in the absence of telomerase (*tert^-/-^*), i.e., occurred at an earlier age than WT. The more refined STEM-based analyses identified 6 specific gene changes accelerated from the age of 9-16 months in *tert^-/-^* and a further 11 from the age of 18-24 months, contrasting to old WT (30-36 months). These included namely down-regulation of SI:DKEY-225F5.4 (an ortholog to ZWINT human gene) and ZGC:101100 (aka dmbx2 in zebrafish and diencephalon/mesencephalon homeobox 2) and upregulation of *klf9* (Kruppel like factor 9, *lyn* (LYN proto-oncogene, Src family tyrosine kinase), mhc1uka (major histocompatibility complex class I UKA) and mxc (myxovirus (influenza virus) resistance C) accelerated from the ages of 9-16 months in *tert^-/-^.* We identified a further 11 from the age of 18-24, contrasting to 30-36 months in WT (**Fig 1C** and SS data). We performed a similar analysis using the broader “DEGs of old age” (**Supp Fig 1C** and SS data). To do this, we compared DEGs identified in *tert^-/-^* zebrafish brains over time (i.e. 2-6 *VS* 9-16 months and 2-6 *VS* 18-14 months) with the WT “DEGs of old age”. We then selected only the ones that changed in *tert^-/-^* in the same direction as WT, i.e., either up or downregulated and further compared these with the corresponding age in sibling WT DEGs, to check whether they were accelerated in *tert^-/-^*, i.e., changed at younger ages than the equivalent sibling WT. Using this strategy, we identified 1049 DEGs of old age accelerated in the absence of telomerase (*tert^-/-^*), which comprised c.36% of the total (2887) “DEGs of old age” (**Supp Fig 2** and SS data). Within these, 4 “DEGs of old age” were accelerated from the early age of 2-6 months in *tert^-/-^*; 639 from the age of 9-16 and 109 from the age of 18-24, contrasting with the age of 30-36 in WT. Notably, when mapped to the ageing transcriptomic hallmarks of ageing, the “DEGs of old age” accelerated in the absence of telomerase identified dysregulation of gene expression, stress response and dysregulation of immune genes as the top 3 significantly enriched transcriptional hallmarks^67^, similarly to the overall DEGs of WT old, and to similar extents (**Supp Fig 3** and SS data).

### 2.2 Telomerase (tert) depletion accelerates ageing-associated stress response in the zebrafish brain

Given our transcriptomic analysis results in 2.1, we proceeded to testing whether these changes were reflected at cell and tissue level starting with markers associated with the “stress response” transcriptomic hallmark of ageing: cell cycle (also highlighted in both the GOBP and the hypergeometric analysis); senescence-associated markers; DNA damage response and relative telomere length, in brain sections of WT and *tert^-/-^* zebrafish over their lifecourse.

There is not a single marker that can be used on its own to irrefutably identify a senescent cell and markers often described to be associated with senescence in proliferative tissues, such as P16 and P21, are highly expressed in terminally differentiated neurons^68, 69^. This may confound identification of putative senescent cells. Nevertheless, markers such as P16 have previously identified as a good marker for predicting disease progression in Parkinson’s Disease (PD) patients^70^, and with ageing^71^. Acknowledging these limitations, we performed a broad characterisation of putative cellular senescence as a stress response using the following **5 main readouts**^42^ (Hernandez-Segura et al., 2018) (**Fig 2B**): **1)** accumulation of Senescence-Associated β Galactosidase (SA-β-Gal) a known biomarker of cellular senescence, at least in most of the cell types tested^13^, and often present in different types of senescence (**Fig 2B1-3**); **2)** levels of expression of cell cycle kinase inhibitors *p21* and *p16* genes (**Fig 2B2, B4 and Supp Fig 4B**); **3)** proliferation, by Proliferating Cell Nuclear Antigen (PCNA) expression and EdU pulse chase labelling (**Fig 2B2, B5 and C**); **4)** DNA Damage Response (DDR) by identifying cells with strong signal reflecting multiple γH2AX foci (**Supp Fig 5**) and **5)** relative telomere length (**Fig 2 D and Supp Fig 5B**).

Our initial whole brain SA-β-Gal staining in WT zebrafish brains detected more intense blue staining in WT old (30-36 months), when compared to young (2-6 months), which motivated us to explore further (**Fig 2A**). To do this, we dissected and sectioned a separate set of brains to determine which regions had increased SA-β-Gal staining, comparing WT and *tert^-/-^* siblings at different ages. Our results showed a significant increase in SA-β-Gal with ageing in specific regions of the WT zebrafish brain, such as in the midline of the telencephalon, diencephalon, cerebellum and medulla oblongata (**Fig 2B1**). Accumulation of SA-β-Gal in the diencephalon, cerebellum and medulla oblongata was significantly accelerated in the absence of telomerase (*tert^-/-^*), where, at 2-6 months, the *tert^-/-^* brain is already displaying the pattern and levels of SA-β-Gal observed in old WT brains of 30-36 months of age (diencephalon quantification shown as a representative example in **Fig 2B2**).

We then collected 2 further sets of zebrafish WT and *tert^-/-^* sibling brains to assess a) *cdkn2a/b* “*p16-like” (RNA in situ),* PCNA (IF) *and* SA-βgal (chromogenic) (**Fig 2**) and b) *cdkn1a* “*p21-like”* and *cdkn2a/b* “*p16-like”* genes (RNA *in situ)* (alongside a repeat of chromogenic SA-βgal as control) (**Supp Fig 4**) in sequential sections of the respective animals. Our data show an increase in *cdkn2a/b* “*p16-like”* expression levels with ageing in WT and *tert^-/-^*, this is not accelerated by the absence of telomerase, either at 2-6 or 9-16 months (**Fig 2B and B4**). Notably, the increase in *cdkn2a/b* “*p16-like”* expression was not accompanied by *cdkn1a* “*p21-like”* expression (**Supp Fig 4B**). We detected very little proliferation in the zebrafish brain—generally under 5% of PCNA+ cells per FOV in any region we looked (**Supp Fig 5A**). Even after 3 days pulse with EdU, the % of EdU^+^ cells in the diencephalon/Optic tectum, where known proliferative niches reside), was under 1%/ FOV (**Fig 2C**). Nevertheless, both PCNA and EdU data show a significant decrease in proliferating cells with ageing in both WT and *tert^-/-^* in the diencephalon/optic tectum (PCNA and EdU), but this is not significantly accelerated in the absence of telomerase at the ages tested.

Another common marker of senescent cells is the accumulation of DNA damage response (DDR) markers, such as γH2AX foci. Our data show that very few cells accumulate γH2AX foci in the aged brain, in all regions tested (never more than 2% of cells (**Supp Fig 5A**). Nevertheless, we detected a significant increase in γH2ax^+^ cells with ageing in both WT and *tert^-/-^* in the cerebellum (**Supp Fig 5A2**) and when we combine diencephalon with Optic Tectum regions (**Fig 2B6**). Notably, accumulation of DDR with ageing is accelerated in the absence of telomerase in these brain regions from the age of 9-16 months in *tert^-/-^* zebrafish. Of note, the senescence-associated markers tested (increase in SA-β-Gal, *cdkn2a/b* “*p16- like”* and γH2AX^+^ cells) occur in both proliferative and non-proliferative areas of the brain (**Fig 2B3)** (see example of ventral diencephalon and dorsal diencephalon, respectively, in a young WT brain (**Fig 2 B3.1**).

To test whether increase in senescence-associated markers correlated with shorter telomeres (i.e. putative replicative senescence), we measured relative telomere length in brain sections of WT fish over time, alongside their *tert^-/-^* counterparts as previously described by us, using a telomeric Peptide Nucleic Acid (PNA) probe alongside a centromeric PNA probe in Fluorescence *in situ* Hybridisation (FISH)^72^ (**Fig 2D** and **Supp Fig 5B**). We show that relative telomere length decreases with ageing in WT brains and this is accelerated in the absence of telomerase, where telomeres are significantly shorter in *tert^-/-^* from the young age of 2-6 months, as has been shown for multiple tissues^20^. However, as expected, relative telomere length in the brain is significantly longer than in the highly proliferative gut epithelia of the same individuals, both in WT and *tert^-/-^* at the ages tested.

Different types of senescence stimuli have been identified to express different SASP factors. We took advantage of available public databases of genes changes associated with SASP factors in humans and mice (SASP atlas and Senmayo databases)^73, 74^ and compared them with our transcriptomic STEM and broad “DEGs of old age”. We were particularly interested in determining whether any of these putative SASP-associated genes were accelerated in the absence of telomerase, at the ages where we see acceleration of other senescence-associated markers. The STEM-based SASP analysis identified 2 genes in common between WT and *tert^-/-^* profiles (*tyms* and *psm2*) (**Supp Fig 6A**), but none of these were accelerated in the absence of tert (Fig 1 D).

However, when we performed the broader “DEGs of old age” accelerated in the absence of telomerase, we identified a further 52 genes, including *psm2 (*accelerated in the absence of telomerase from the age of 9-16 months), which was also detected in the STEM profiles (**Supp Fig 6B**). Of note, *idh1, lancl1, masp1, ncl and tuba1a* are accelerated in the absence of telomerase from the age of 18-24 months, and the remaining ones from 18-24 months in *tert^-/-^* (SS data).

### 2.3 Telomerase (tert) depletion accelerates ageing-associated inflammation in the zebrafish brain

We then proceeded to testing whether the changes associated with the “dysregulation of immune response” ageing transcriptomic hallmark—also highlighted in both the GOBP and hypergeometric analysis—were reflected at cell and tissue level starting. We started by quantifying the percentage of immune cells in in brain sections of WT and *tert^-/-^* zebrafish over their lifecourse.

Because there is no agreed specific microglia marker in zebrafish, we combined the tg(*mpeg*1.1-mCherry-caax) transgenic zebrafish line^75^, which has been shown to identify macrophages and microglia in the brain^76^, with a well-described pan-immune cell marker (L-plastin or LCP-1)^77–80^ and use immunofluorescence (IF) to quantify immune cell populations in the brain^81^ as a readout for inflammation. Using these two markers allowed us to identify macrophages/microglia as L-plastin^+^*mpeg*^+^ cells, and distinguish them from any other immune cells, which would be L-plastin^+^*mpeg*^-^ (**Fig 3A).** Our data show that there is an increased number of plastin^+^*mpeg*^+^ cells (**Fig 3A1**) but not of L-plastin^+^*mpeg*^-^ cells (**Fig 3A2**) with ageing, in most of the zebrafish brain (WT 2-6 months *VS* WT 30-36 months) (**Supp Fig 7A**), suggesting a macrophage/microglia specific increase. The macrophage/microglia increase in the brain with ageing (WT 2-6 months *VS* WT 30-36 months) is suggestive of neuroinflammation and was accelerated in the absence of telomerase from the early age of 2-6 months (WT 2-6 months *VS tert^-/-^* 2-6 months), at which ages we do not see any significant increase in immune cell proliferation (**Supp Fig 7B**).

**Figure 3.**
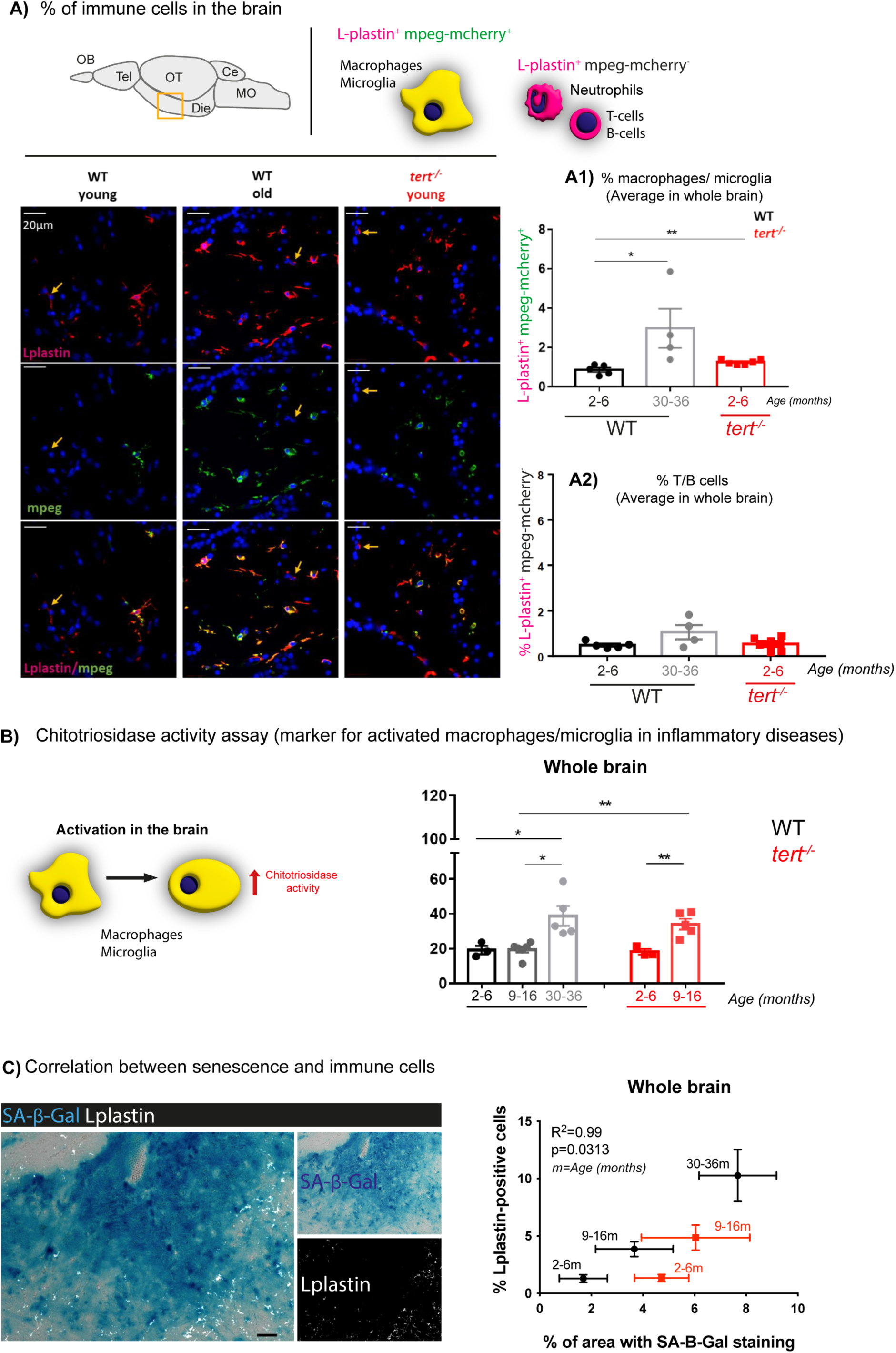
Telomerase depletion is associated with increased inflammation in the aged zebrafish brain. (A) Representative images of mpeg-mcherry and L-plastin staining in the diencephalon of adult zebrafish (schematic figure highlights the region imaged as diencephalon). The yellow arrows highlight Lplastin^+^; mpeg^-^ cells. (A1) Quantifications of L-plastin-positive; mpeg-positive cells (orange; from red and green co-localisation) and (A2) L-plastin-positive; mpeg-negative cells (red) in the whole zebrafish brain show an increased number of macrophages/microglia (L-plastin-positive; mpeg-positive cells) with natural ageing, and this is accelerated in the absence of telomerase (in *tert^-/-^* at 2-6 months of age). However, we observed no differences in the number of T/B cells and neutrophils (L-plastin-positive; mpeg-negative cells) with ageing, in the WT or *tert^-/-^* fish. N=-6 per group. (B) Schematic figure of the chitotriosidase assay (left). Quantifications (right) show that chitotriosidase activity increases with natural ageing at >30 months of age, and that this is accelerated in the *tert^-/-^* at the age of 9-16 months. (C) Representative images of co-staining with SA-β-Gal (blue) and L-plastin (white) in brain sections show that most of the L-plastin-positive cells do not co-localise with the SA-β-Gal staining (left). Staining quantification shows a significant correlation between increased number of immune cells and increased expression of SA-β-Gal with ageing in both WT and *tert^-/-^* (right). (A1, A2, B) Each dot represents one animal. (A1, A2, B, C) Bar errors represent the SEM. * <0.05; ** <0.01; *** <0.001.

To understand whether macrophage/microglia increase with ageing was associated with immune activation, we used a chitotriosidase activity assay. Chitotriosidase is an enzyme expressed by activated macrophages and other phagocytic cells and can therefore be used as a biomarker of microglia/macrophage activation, including in zebrafish (Keatinge et al., 2015). Our data show that there are increased levels of chitotriosidase in the naturally aged brain (WT 2-6 *VS* 9-16 *VS* 30-36 months), suggesting age-associated neuronal immune activation, which we show is accelerated in the absence of telomerase (*tert^-/-^*) from the age of 9-16 months (**Fig. 3B**).

Accumulation of cellular senescence is thought to contribute to chronic inflammation in ageing, often termed “inflammageing”^82–84^. On one hand, in some contexts, immune cells have been shown to be recruited to sites of cellular senescence^85, 86^. On the other hand, senescent cells have been shown to release pro-inflammatory factors (part of SASP), which can contribute to paracrine senescence^87^ as well as to tissue inflammation^83^. We therefore set out to determine how the increased levels of senescence-associated markers we detected in the zebrafish brain correlated with immune cell numbers. To do this, we performed IF staining with the pan-immune cell marker L-plastin on SA-β-Gal-stained brain tissue sections. Even though we cannot exclude that some of these immune cells may also be SA-β-Gal^+^ themselves, most of the immune cells appear to surround, rather than overlap with SA-β-Gal^+^ areas (**Fig 3C)** and there is a statistically significant positive correlation between % of SA-β-Gal^+^ areas and % of immune cells with ageing in the WT zebrafish brain (R^2^=0.99), p(two-tailed) =0.0313.

### 2.4 Telomerase (tert) depletion accelerates ageing-associated blood-brain barrier dysfunction

Because our data show that telomerase (tert) depletion accelerates ageing-associated neuroinflammation (**Fig 3**) not explained by increased immune proliferation (**Supp Fig 7B**), we hypothesised that there could be immune infiltration from the periphery. Immune cells should not cross the Blood-brain barrier (BBB) and invade the brain in a healthy condition unless the BBB is compromised (reviewed in^88^). It has been described that zebrafish have a functional BBB from the 10th day post fertilisation (dpf)^89^ after which molecules such as horseradish peroxidase (HRP, 44,000Da), evans blue (961Da), and sodium fluorescein (376Da) do not cross this barrier^89, 90^.

To test whether zebrafish BBB becomes leaky with ageing and whether telomerase plays a role in it, we set up a BBB permeability assay in both WT and *tert^-/-^* at similar ages where increased immune activation (chitotriosidase activity) was identified in the brain. To do this, we performed intraperitoneal injections (IP) of an infrared fluorescent dye IRDye 680RD Dextran diluted in saline (Hanks buffered solution-HBSS). We confirmed this dye’s suitability for assessing BBB by performing a pilot study comparing dispersion of conjugated molecules of different sizes throughout the body at different time points and determined that 4kDa Dextran conjugated with fluorescent dye was optimal for assessment of tissue dispersal 3 hrs post-injection (not shown). Three hours after injection, fish were culled, and dissected brain fluorescence was measured (using a Li-cor Odyssey imaging system). Our data show that BBB permeability increases with WT ageing, as measured by 4kDa dextran-FITC-associated fluorescence in the brain, which was accelerated in the absence of telomerase (*tert^-/-^*) from the tested age of 9-16 months.

**Figure 4.**
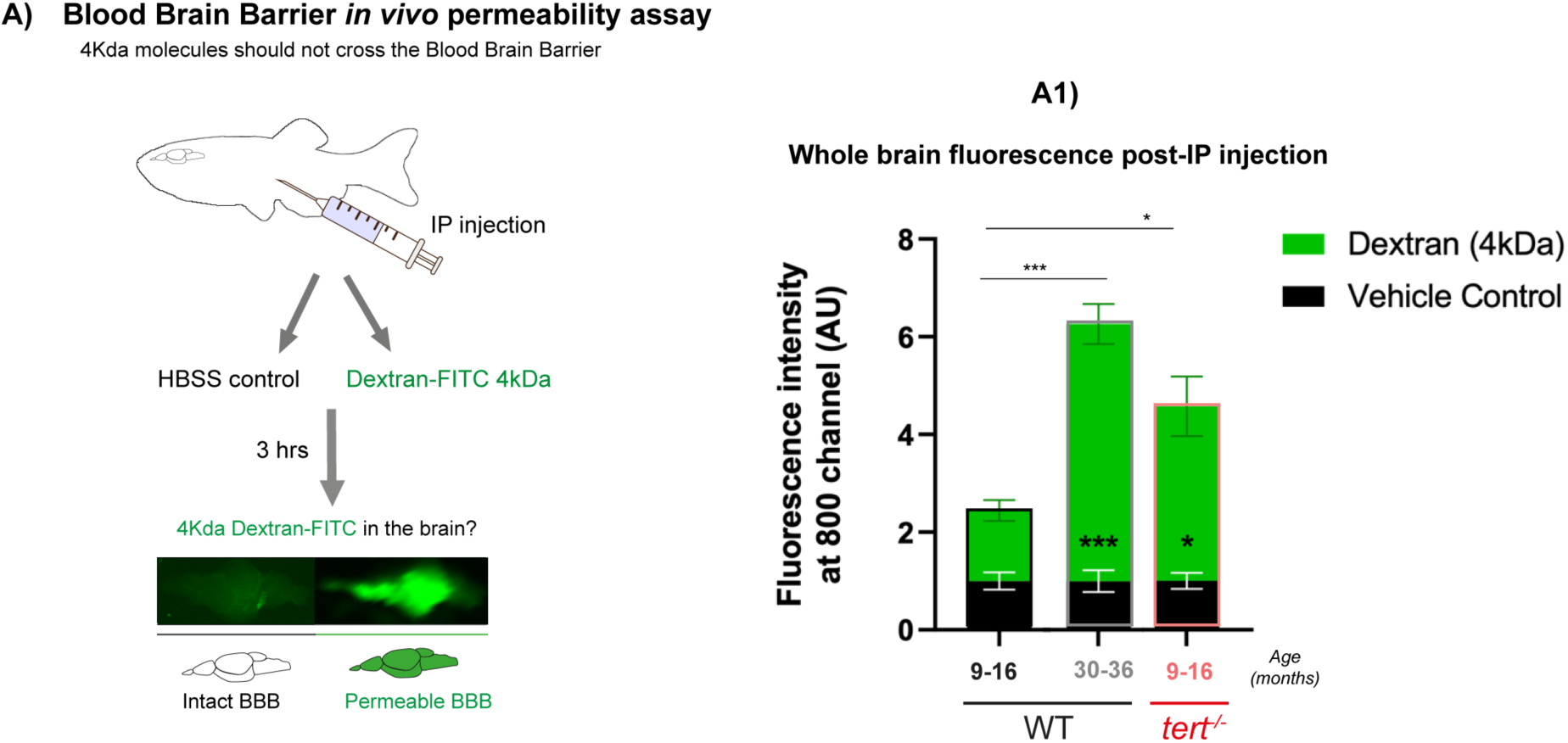
Telomerase depletion is associated with increased BBB permeability with ageing in the zebrafish brain. (A) Schematic figure of the BBB permeability assay. Dextran-FITC 4kDa or vehicle control (HBSS) were injected by intraperitoneal injection. 3 hours later the brains were dissected and imaged in the Odyssey® CLx (Li-cor) to confirm the presence of Dextran-FITC. Quantification of the green fluorescence in the fish brains shows higher FITC expression in old WT (>30 months) compared to middle aged ones (9-16 months). This increased presence of dextran-FITC in *tert^-/-^* is already observed at 9-16 months, overall suggesting that there is increased brain permeability with natural ageing and that this is anticipated in the absence of telomerase. N=9-20 per group in the tested animals; N=3-6 per group in the vehicle control animals. Bar errors represent the SEM. Statistics: unpaired t-tests. * p<0.05; ** p<0.01; *** p<0.001.

### 2.5 Telomerase (tert) depletion accelerates ageing-associated behavioural changes in zebrafish

It is known that inflammation can influence behaviour. For example, lipopolysaccharide (LPS)-induced peripheral inflammation has been shown to lead to behavioural alterations in rodents, in a dose-dependent manner, including changes in anxiety-like behaviour^91, 92^. Given that we observed a telomerase-dependent increase in neuroinflammation with ageing, we set out to test whether it associated with any alterations in behaviour. To do this, we assessed anxiety-like behaviour using a novel tank test ^93, 94^ in WT and *tert^-/-^* zebrafish at young and old ages (**Fig 5A**). The novel tank test showed that, with ageing, WT fish tend to spend less time in the top part of the tank, particularly at >30 months of age, indicative of increased anxiety-like behaviour. This behaviour is already detected in *tert^-/-^* at the earlier age of 9-12 months (**Fig 5B**). Importantly, no differences between groups were observed in the total distance swam throughout the test (**Fig 5C**). Therefore, the differences observed in the time spent swimming in the top area of the tank, in advanced ages, is likely to be associated to an anxiety-like behaviour phenotype rather than to potential locomotive defects. Together, lower time spent in the top part of the tank suggests reduced anxiety-like behaviour with natural ageing, which our data suggest is a telomerase-dependent phenotype of old age.

**Figure 5.**
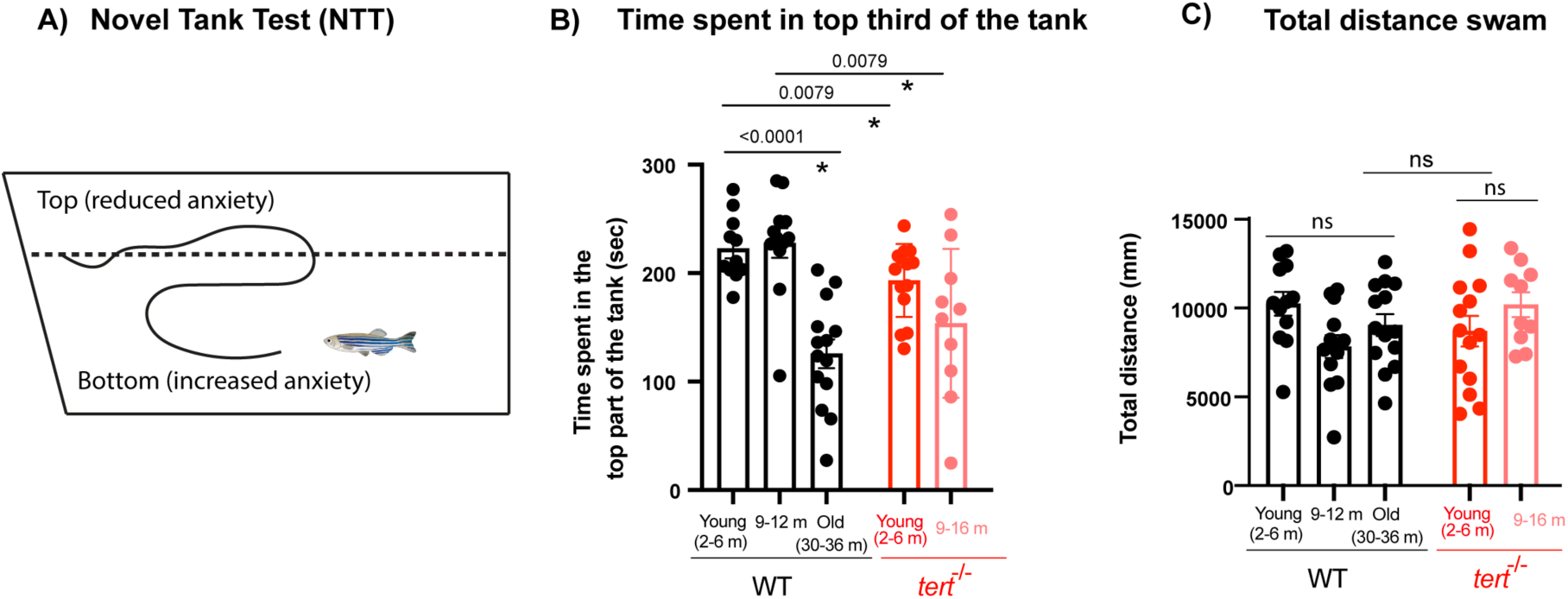
Telomerase depletion is associated with increased anxiety-like behaviour. (A) Schematic figure of the setup and behaviour assessment during the novel tank test (NTT). (B) Quantifications show that old WT (>30 months) spend less periods of time in the top part of the tank compared to younger siblings, and that this behaviour is already observed in *tert^-/-^* at 9-16 months. (C) Quantifications of the total distance swam during the entirety of the test show no differences between groups, suggesting that the differences observed in the time spent in the top area of the tank are not explained by a locomotive defect in advanced ages. N=10-14. Each dot represents a fish. Bar errors represent the SEM. * p<0.05; ** p<0.01; *** p<0.001.

## 3 DISCUSSION

Decreased telomerase expression, telomere shortening, senescence-associated markers and inflammation have all been independently observed in the ageing brain and associated with human neurodegeneration. However, causality between limited telomerase expression and increased brain senescence and neuro-inflammation in a WT setting is yet to be established. Here, we address these questions using the zebrafish as an ageing model which, akin to humans, displays premature ageing and death in the absence of telomerase and where telomere shortening is a driver of cellular senescence. We hypothesised that limited expression of Tert in the aged zebrafish brain contributes to brain degeneration with ageing. To our knowledge, this manuscript represents the first broad characterisation of zebrafish brain ageing in the presence and absence of telomerase (Tert).

Our transcriptomic analyses highlighted increased inflammation and decreased cell cycle as key signatures of ageing accelerated in the absence of telomerase (tert) which were reflected at cell/tissue level and accompanied by behaviour alterations. Interestingly, some ageing-associated phenotypes were accelerated from as early as 2-6 months in the *tert^-/-^*, whereas the majority were from 9-16 months, which led us to our current working model (Fig 6). In specific, telomerase depletion accelerates ageing-associated telomere shortening, accumulation of SA-β-Gal, a lysosomal protein, which are generally known to accumulate with senescence; and microglia/macrophages numbers from the young age of 2-6 months in *tert^-/-^*.These tissue-level changes are accompanied by an increase of ageing-associated increase in anxiety-like behaviour from that same early age in *tert^-/-^.* Our data leads us to suggests a potential model in which SA-βGAL accumulation and recruitment of immune cells. Whether this increased neuroinflammation is detrimental remains to be determined, and it is not entirely clear in human neurodegeneration. Neuroinflammation is associated with neurodegeneration and disease in humans^50, 95, 96^, but causality is not clear. It is not clear from our data whether neuroinflammation in the aged zebrafish brain is contributing to the senescence phenotypes via paracrine senescence mechanisms^87^ or whether immune cells are “trying” to clear senescent cells^97^. Nevertheless, even if initially recruited for clearance purposes, this function may be decreased with ageing and thus chronic inflammation may contribute to a senescence phenotype by increasing senescence-associated markers such as SASP and DDR. Our data specifically highlights *psme2* (proteasome activator subunit 2), as a particularly relevant putative SASP-associated factor, since it was identified in both our STEM and broader DEGs of old age accelerated in the absence of telomerase (**Supp Fig 6**). loss of proteostasis is a known hallmark of ageing and *psme2* may be involved in this crosstalk^5^, Potentially as a consequence of increased inflammation and accumulation of cellular senescence, our data show that telomerase depletion accelerates ageing-associated increase in immune activation (increased chitotriosidase activity) and BBB permeability. In turn, BBB leakiness is likely to lead to further recruitment of immune cells from the periphery, further exacerbating inflammation and BBB dysfunction in a positive feedback loop. Of note, our broad analysis of “DEGs of old age” showed that *ctss2.1* is significantly upregulated with ageing, which has recently been implicated in macrophage-driven age-dependent disruption of the blood-CSF barrier ^98^. BBB leakiness is likely to lead to further recruitment of immune cells from the periphery potentially exacerbating inflammation and BBB dysfunction in a positive feedback loop.

**Figure 6.**
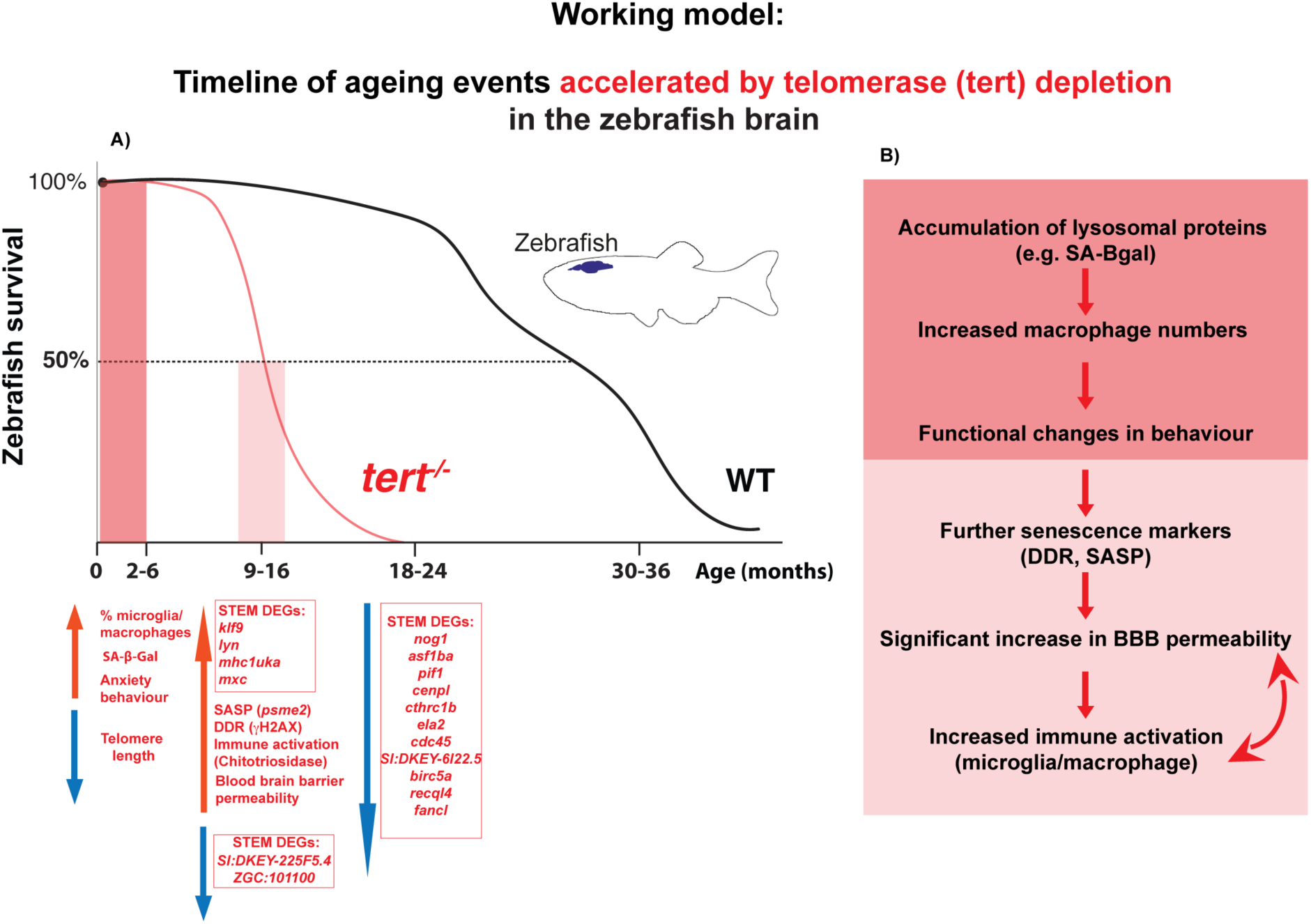
Working model: timeline of ageing events accelerated by telomerase depletion in the zebrafish brain. (A) Survival curve showing the early to late stages of ageing in WT fish compared to *tert^-/-^* fish, highlighting the changes we identify at cellular, tissue and functional levels with ageing and that are accelerated in the absence of telomerase, at different ages throughout their lifecourse. (B) Table summarising the sequence of events we observe in the ageing brain and that are accelerated by telomerase depletion: Accumulation of lysosomal proteins (such as SA-β-Gal), increased number of macrophages and increased anxiety-like behaviour are already observed at the young age of 2-6 in *tert^-/-^* fish (but only at later ages in WT fish). Dysfunction at lysosomes and recruitment of immune cells are likely to contribute to a senescence phenotype by increasing senescence-associated markers such as SASP and DDR, which are observed at 9-16 months in *tert^-/-^*. This is accompanied by increased immune activation (increased chitotriosidase activity) and BBB permeability, at the same age, potentially as a consequence of increased inflammation and accumulation of cellular senescence. In turn, BBB leakiness is likely to lead to further recruitment of immune cells from the periphery, further exacerbating inflammation and BBB dysfunction in a positive feedback loop.

Together, our work suggests that telomerase has a protective role in the zebrafish brain against the accumulation of senescence-associated markers and neuro-inflammation and is required for BBB integrity. Importantly, the acceleration of senescence-associated markers in the absence of tert occurs not only in the expected proliferative areas but also in non-proliferative ones, where it is unlikely due to telomere-dependent replicative exhaustion, suggesting that non-canonical roles of telomerase may be involved.

## 4 MATERIALS AND METHODS

### 1. Animal husbandry, ethics, genotypes and ages

Zebrafish were maintained at the standard conditions of 27-28°C, in a 14:10 hour light-dark cycle, and fed twice a day with Artemia (live rotifers) and Sparus (dry food). All the experiments were performed in the University of Sheffield. All animal work was approved by local animal review boards, including the Local Ethical Review Committee at the University of Sheffield (performed according to the protocols of Project Licences 70/8681 and PP4455093). Three strains of adult zebrafish (Danio rerio) were used for these studies: wild-type (WT; AB strain), tert^-/ (tertAB/hu34^^30)^ and *Tg(mpeg1:mCherryCAAX)sh378* (Source: University of Sheffield). Wild-type fish were obtained from the Zebrafish International Resource Center (ZIRC). The telomerase mutant line tert ^AB/hu34^^30^ was generated by N-ethyl-nitrosourea mutagenesis (Utrecht University, Netherlands (Wienholds et al., 2003)). It has a T→A point mutation in the tert gene and is available at the ZFIN repository, ZFIN ID: ZDB-GENO-100412–50, from ZIRC. The fish used in this study are direct descendants of the ones used in the first studies describing this line (Carneiro et al., 2016; Henriques et al., 2013), by which point it had been subsequently outcrossed five times with WT AB for clearing of potential background mutations derived from the random ENU mutagenesis from which this line was originated.

The tert ^hu34^^30/hu34^^30^ homozygous mutant is referred to in the paper as *tert^−/−^* and was obtained by in-crossing our tert heterozygous AB/hu3430 strain, and WT and tert-/- fish compared in this study are siblings, derived from that F1 *tert^+/-^* in-cross, following gold standard guidelines to use this line, as recently reviewed here (Henriques & Ferreira, 2024).Similarly, in order to produce WT and *tert^−/−^* siblings carrying the *Tg(mpeg1:mCherryCAAX)sh378* transgene, tert^+/-^; *mpeg1:mCherryCAAX* were incrossed. In this study, we used only males, given that the kinetics of telomerase-dependent ageing in zebrafish has been well described for males only^18, 20^, which is a limitation of the model and this study. Nevertheless, there is data showing “Limited sex-biased neural gene expression patterns across strains in Zebrafish”^99^, giving strength to our study.

Genotyping was performed by PCR of the tert gene as before. To study age-related phenotypes in the zebrafish brain, we used 30-36 months old fish for what we consider old in WT (in the last 25-30% of their lifespan), and we considered the *tert^-/-^* old fish at the equivalent age (18-24 months), which approximately corresponds to the last 25-30% of their lifespan. In specific, ‘Old’ was defined as the age at which most of the fish present age-associated phenotypes, such as cachexia, loss of body mass and curvature of the spine. These phenotypes develop close to the time of death and are observed at 30-36 months of age in WT and at >18-24 months in *tert^-/-^*^18, 20^.

#### Note on Zebrafish gene and protein nomenclature

This manuscript followed the agreed convention as described in: https://zfin.atlassian.net/wiki/spaces/general/pages/1818394635/ZFIN+Zebrafish+Nomenclature+Conventions

#### In summary

##### Gene nomenclature

Genes should be named after the mammalian orthologue whenever possible. When mammalian orthologues are known, the same name and abbreviation should be used, except all letters are italicized and lower case. Members of a gene family are sequentially numbered.

##### Examples

Names - engrailed 1a, engrailed 2b Symbols - eng1a, eng2b

##### Protein nomenclature

The protein symbol is the same as the gene symbol, but non-italic and the first letter is uppercase.

Note the differences between zebrafish and mammalian naming conventions:

##### species / gene / protein

zebrafish /*shha*/ Shha

human / *SHH* / SHH

mouse / *Shh* / SHH

### 2. Tissue fixation

<u>Paraffin-embedded sections</u>: Fish were culled by immersion in concentrated tricaine (MS-222). After terminal anaesthesia, the animals were fixed by immersion in 50 ml of Neutral buffered formalin (NBT, VWR International, Radnor, PA, USA) at 4°C for 48h-72h and decalcified in 50 ml of 0.5M ethylenediaminetetraacetic acid (EDTA), pH 8, for additional 48-72h. Then, the whole fish were paraffin-embedded (in formalin I for 10min, formalin II for 50min, ethanol 50% for 1h, ethanol 70% for 1h, ethanol 95% for 1h 30min, ethanol 100% for 2h, ethanol 100% for 2h 30min, 50%:50% of ethanol 100% : Xilol for 1h 30min, xylene I for 3h, Xylene II for 3h, paraffin I for 3h, paraffin II for 4.30h), longitudinally sliced (sagittal slices, 3 μm thick, using the Leica TP 1020) and stored at room temperature (RT). To do this, diluted ethanol absolute (VWR International) and paraffin histosec pastilles (Merck & Co, Kenilworth, NJ, USA) were used.

<u>Cryosections</u>: Fish were culled by immersion in concentrated MS-222, followed by immediate decapitation (Schedule 1 method). The brain was dissected, transferred to and rinsed in cold L15 (Thermo Fisher Scientific, Waltham, MA, USA). Afterwards, the brain was fixed in 4ml NBT overnight at 4°C, washed in 3ml cold Phosphate Buffered Saline (PBS, 1x), and immersed in 4ml of 30% sucrose in PBS overnight at 4°C for cryoprotection. Finally, the tissue was mounted in a base mould (Electron Microscopy Sciences, Hatfield, PA, USA), with O.C.T. (VWR), snap frozen, and stored at -20°C. Prior to use, 13μm coronal tissue sections were cut using a Leica CM1860 cryostat, attached to SuperfrostPlus slides (Epredia) and air dried for a minimum of 2h at RT. Slides were stored at -20°C for a maximum of 3 months.

### 3. RNA probe synthesis for *in situ* hybridisation (ISH)

A Zebrafish p16-like fragment was amplified using the following primers: FW cgttgaacATGAACGTCG and REV tctttaaacattTTAAACACGATTGAG and cloned into the pCR-Blunt ii-TOPO vector (Thermofisher Scientific Inc, USA), For anti-sense probe generation, the plasmid was linearised with HindIII (New England Biolabs) and synthesis carried out with T7 RNA polymerase (New England Biolabs) and DIG-labelling mix (Merck Life Science UK Limited).

For the sense probe, plasmid was linearisated with XhoI (New England Biolabs), and probe synthesised with SP6 RNA polymerase (New England Biolabs).

Probes were cleaned up using SigmaSpin post-reaction clean-up columns (Sigma Aldrich) and used at a final concentration of 3ng/μl for hybridisation.

### 4. Chromogenic RNA fluorescence *in situ* hybridisation (RNA-FISH)

Cryosections were fixed with NBT for 10min at RT followed by 3x washes with Diethyl pyrocarbonate (DEPC)-treated PBS. Sections were then acetylated with 11.2 μl triethanolamine and 2.5 μl acetic anhydride per 1ml H_2_O for 10 min at RT (both Sigma Aldrich). After 3x washes in DEPC-PBS, sections were incubated in prehybridization solution (prehyb) and coverslipped (prehyb: 50% formamide, 5x SSC pH7, 2% Blocking reagent, 0.1% TritonX100, 0.5% CHAPS, 1mg/ml yeast RNA, 50 μg/ml heparin sodium salt (all from Sigma Aldrich)) for 1h at 68°C in a humidified box. The solution was replaced with 200μl l fresh prehyb containing 3ng/ μl DIG-labelled RNA probe, coverslipped and incubated overnight at 68°C in a humidified box.

Following this, coverslips were removed and slides were washed for 1h in prewarmed wash #1 (50% formamide, 5xSSC pH4.5, 1%SDS) at 68°C, then 1h in prewarmed wash #2 (50% formamide, 2xSSCpH4.5, 1% Tween20) at 68°C in coplin jars.

Slides were washed in TBS-T (8g NaCl, 0.2g KCl, 25ml 1M Tris pH7.5, 10ml tween20 per L solution) at RT, blocked in TBS-T/10% Heat inactivated goat serum for 40 min at RT, then incubated in fresh block/alkaline phosphatase-conjugated anti-DIG at 1:2000 for 80min at RT. Slides were washed extensively in TBS-T followed by 3x washes in freshly prepared NTMT (for 50ml solution:1ml 5M NaCl, 2.5ml 2M Tris pH9.5, 2.5ml 1M MgCl2, 500ul Tween20, 43.5ml ddH2O). Development was started by incubation in filtered staining solution ((4.5ul NBT +3.5ul BCIP per 1ml NTMT (both Merck Life Science UK Limited)) at RT in the dark. Development was monitored daily with replenishment of fresh filtered staining solution. Once the signal was clear, with minimal background staining, the reaction was stopped by washing in PBS. Slides were mounted in glycergel (Agilent, Santa Clara, CA, USA) or vectashield (2BScientific) before imaging on a Nikon Ti inverted microscope at 10x. Images were tiled using Nikon elements software. For analysis, the diencephalon and optic tectum (including PGZ) were delineated and thresholded using fiji image J to determine the area of positive staining within each macroarea. The percentage of positive tissue per section macroarea was then determined.

### 5. Immunofluorescence (IF)

Paraffin sections with whole fish slices were dewaxed and hydrated through the following sequence of washes: histoclear (Scientific Laboratory Supplies, Wilford, Nottingham, UK, 2x5 min), 100% ethanol (Thermo Fisher Scientific, 2x5 min), 90% ethanol (5 min), 70% ethanol (5 min), and distilled water (2x5 min). Cryosections were hydrated in PBS for 10 min before proceeding. After antigen retrieval in 0.01M citrate buffer at pH 6.0 (sub-boiling at 800 W in a microwave for 10 min), the tissue sections were permeabilized in PBS 0.5% Triton X-100 (for 10 min) and blocked in 3% bovine serum albumin (BSA), 5% Goat Serum in 0.3% Tween-20 in PBS (for 1 h). Then, primary antibody incubation (overnight at 4°C) was followed by washes in PBS 0.1% Tween-20 and secondary antibody incubation (1 h at RT or overnight at 4°C). Finally, the tissues were incubated in DAPI for nuclear staining (Thermo Fisher Scientific), 1:10,000 dilution in PBS from a 10mg/ml stock, for 10 min; washed in PBS 1x, mounted with vectashield (Vector Laboratories, Burlingame, CA, USA), covered with a coverslip (Scientific Laboratory Supplies), and sealed with clear nail varnish. For details on the primary and secondary antibodies refer to Table 1 and Table 2, respectively.

**Table 1.**
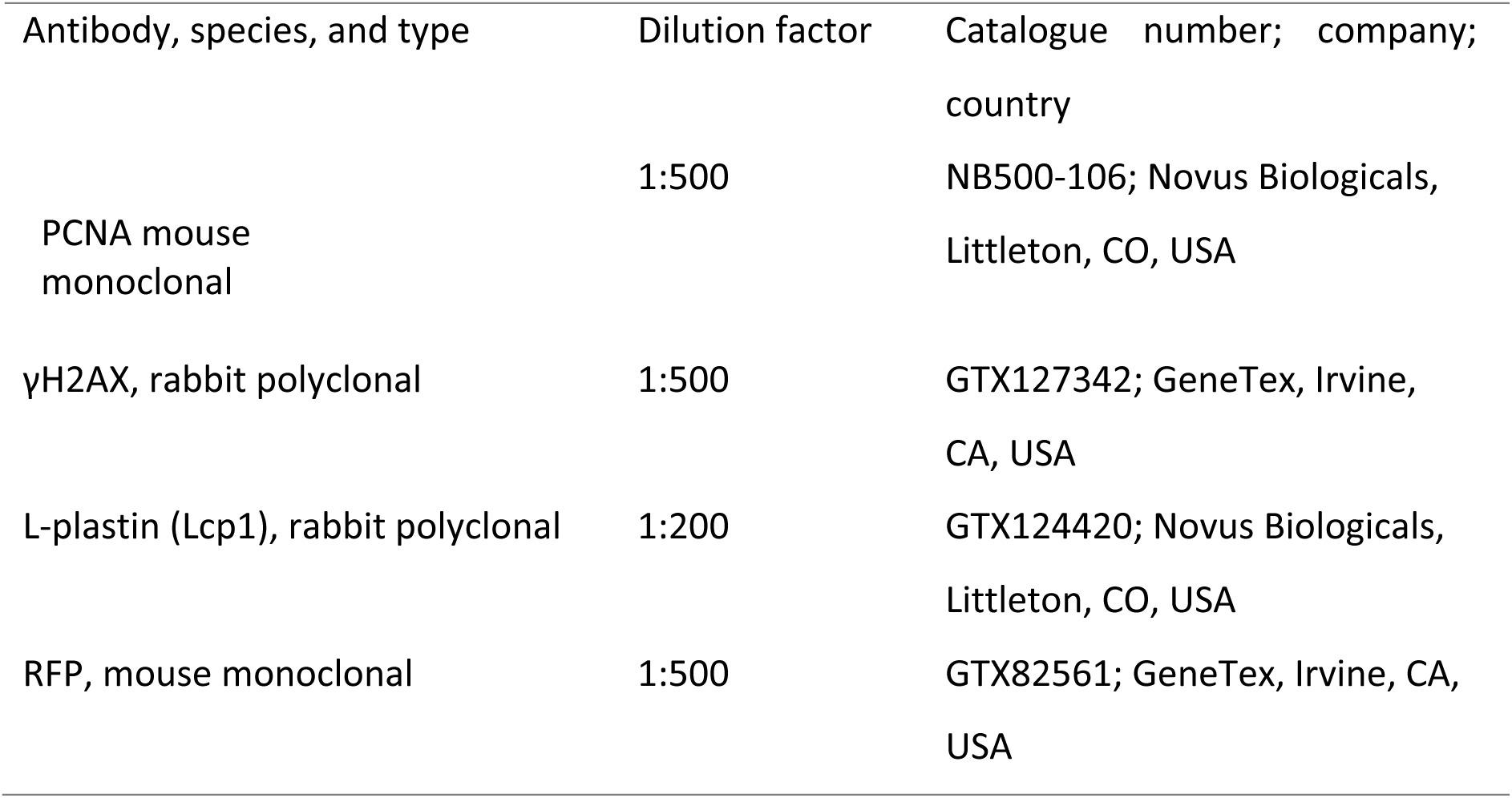
Primary antibodies used for immunostaining.

**Table 2.** Secondary antibodies used for immunostaining.

| Antibody, species, and type | Dilution factor | Catalogue number; company; country |
| --- | --- | --- |
| Goat anti-rabbit IgG Alexa Fluor<sup>®</sup> 488 | 1:500 | A11008; Invitrogen, Carlsbad, CA, USA |
| Goat anti-rabbit IgG Alexa Fluor<sup>®</sup> 568 | 1:500 | 10032302; Thermo Fisher Scientific, Waltham, MA, USA |
| Goat anti-mouse IgG Alexa Fluor<sup>®</sup> 647 | 1:500 | A21235; Thermo Fisher Scientific, Waltham, MA, USA |

### 6. Telomere-fluorescence *in situ* hybridization (telo-FISH)

Telo-FISH was performed after IF. Briefly, after incubation with the secondary antibody and respective washes, the tissues were fixed in 4% PFA (SIGMA-ALDRICH) for 20min. Washes in PBS, 3x 10min were followed by a dehydration step consisting of washes with ice cold 70%, 90% and 100% ethanol (Thermo Fisher Scientific, 3min each). After this, the slides were left to dry for 1h, 10μl of hybridisation solution was pipetted on top of each tissue section and the slides were covered with a coverslip. Hybridisation solution (for a total volume of 100μl): 8.56μl MgCl2 buffer pH 7.0, and 82mM sodium hydrogen phosphate (SIGMA-ALDRICH), 70μl of deionised formamide (Thermo Fisher Scientific), 1μl of 1mM Tris, pH 7.5 (SIGMA-ALDRICH), 5μl of 1x blocking buffer (10x blocking reagent (Merck & Co.) diluted to 1x in maleic acid (SIGMA-ALDRICH)), 2μl of PNA probe (Telo-Cy3, zebrafish, 568; Cambridge Research Biochemicals, Cleveland, UK), 2μl of Cent-Fam probe (488; Eurogentec, Liège, Belgium), and 11.44μl of dH2O.

The slides were incubated for 10min at 80°C for DNA denaturation, followed by incubation for 2h at RT, in the dark, to allow the probe to hybridise with the complementary DNA. After the incubation period, coverslips were gently removed and the slides were washed for 10min in a solution made of 70% formamide (SIGMA-ALDRICH), 10% 2x saline-sodium citrate (SSC, SIGMA-ALDRICH) and 20% dH2O. This was followed by two more washes (10min each), in 2x SSC. SSC was diluted from a 20x stock solution consisting of 88.3g of Tri-sodium citrate dihydrate (SIGMA-ALDRICH) and 175.3g of NaCl (SIGMA-ALDRICH), diluted in dH2O to a total volume of 1L, adjusted to pH of 7.0 and autoclaved. Finally, the slides were washed in PBS, incubated in DAPI (Thermo Fisher Scientific) for 10min and mounted as previously described in section 2.3.2.

### 7. IF and telo-FISH imaging and quantification

IF and telo-FISH staining was imaged using the DeltaVision microscope, Nikon Widefield inverted microscope or a Nikon Ti2 W1 spinning disc confocal at the Wolfson Light Microscopy Facility (LMF), using a 40x objective. An average of 20 images were acquired from the brain of each animal (paraffin sections) for longitudinal paraffin sections and 18 for transverse cryosections (minimum 6 images per macro-area). For images showing whole brain transverse cryosections, either 4x or 10x images were taken, with tiling in Nikon elements software for the latter. Imaging settings were kept consistent within experiments for accurate comparisons.

After acquisition, the images were analysed by manual counting: (1) the number of cells with more than 5 foci of γH2AX (considered γH2AX+ cells) PCNA-, (2) the number of PCNA+ cells, (3) the number of L-plastin^+^, (4) the number of mpeg^+^ cells, and (5) the total number of cells. Telo-FISH staining was quantified by the intensity of the mean of the nucleus. The ratio ‘telomeres/centromeres’ was calculated by dividing the intensity of the mean of telomeres (568 channel) by the intensity of the mean of centromeres (488 channel), which normalises the telomeric probe signal by the centromeric probe signal within individual cells. The brain commonly exhibits cytoplasmic autofluorescence from vascular and other cell types. These were excluded from our analyses by comparison to neighbouring control sections stained only with secondary antibodies. Nuclear markers such as γH2AX, PCNA were only classed as positive when they showed signal in the nucleus as determined by co-labelling with DAPI. For telomeric and centromeric labelling analysis, individual nuclei were circumscribed based on DAPI labelling and mean gray fluorescence determined. Mpeg^+^ and Lplastin^+^ cells were manually counted and could be easily distinguished from autofluorescent cells based on strength of signal and morphology.

The analysis was performed on raw images, using the ImageJ Fiji (v. 1.51). Images were then processed with Abode Illustrator 21.0.2 for display purposes.

### 8. SA-β-Gal

The Senescence-Associated β-Galactosidase Staining Kit (Cell Signalling Technology, Danvers, MA, USA) was used to perform the SA-β-Gal staining, following manufacturer’s instructions.

<u>Whole brain SA-β-Gal staining</u>: Fish were culled with neat Tricaine and decapitated. Whole heads were fixed in 4ml PFA overnight at 4c with shaking. The following day, whole heads were washed 3 x for 1h in PBS at RT with slow shaking. Whole brains were then dissected in fresh PBS and immersed in 2ml in SA-β-Gal staining solution mix, consisting in 1x staining solution (pH 5.9), 1x solution A, 1x Solution B, 1mg/ml Xgal (SA-β-Gal staining kit #9860 by Cell Signaling Technology). and incubated at 37°C overnight with slow shaking in the dark. The following day, brains were washed 3 x for 1h in PBS at RT with slow shaking and imaged using a Nikon SMZ1500 Stereomicroscope (2000).

<u>SA-β-Gal staining on cryosections</u>: Briefly, after rinsing the cryosections in 1x PBS for 5 min, 200ul of β-Gal staining solution was added per slide as above. Then, the slides were coverslipped and incubated at 37°C overnight in a humidified chamber box. The following day, cryosections were rinsed in 1x PBS (3x1min), and mounted in glycergel (Agilent, Santa Clara, CA, USA). Slides were then imaged using an Olympus BX60 microscope or Nikon Ti inverted microscope at 10x with tiling using Nikon elements software.

Image analysis was done using ImageJ Fiji as described for RNA-FISH. Images were then processed with Abode Illustrator 21.0.2 for display purposes.

For SA-β-Gal combined with IF, sections were first labelled with SA-β-Gal as described above, then fixed for 10min with NBT at RT, rinsed and processed for IF as described in methods section 5.

### 9. RNA Extraction

Fish were culled by Schedule 1 method (as described before in section 2.2). Zebrafish brains were dissected, transferred to a microcentrifuge tube with 100 μl of Trizol (Thermo Fisher Scientific), snap frozen in dry ice and stored at -80°C until RNA extraction.

For RNA extraction, the tissue was homogenised with a mechanical homogenizer (VWR International) and 1.5 pestle (Kimble Chase, Vineland, NJ, USA), and an extra 50 μl of fresh Trizol (Thermo Fisher Scientific) was added. After 5 min of incubation at RT, 30 μl of chloroform (1:5, VWR International) was added. The samples were then incubated for further 3 min at RT and centrifuged (at 13,000g for 30 min at 4°C). The resulting upper aqueous phase was transferred to a new microcentrifuge tube, and isopropanol (Thermo Fisher Scientific) was added (approximately 83% of the amount of aqueous phase). After an incubation of 10 min at RT, the samples were centrifuged (13,000g for 15 min at 4°C) and the supernatant was discarded. The pellets were washed in 250 μl of 75% ethanol twice and then left to air dry. In the end, the pellets were resuspended in 14 μl of nuclease-free water (VWR International) and stored at -80°C. All the samples were analysed by Nanodrop 1000 Spectrophotometer to measure the concentration of RNA. Some samples (randomly chosen) were also analysed in the Bioanalyzer to assess RNA integrity. All the samples included in this study had an RNA integrity number (RIN) of >9.

### 10. RNA sequencing

RNA-Sequencing was performed at the Genomics and Sequencing facility of Sheffield Institute for Translational Neuroscience (SITraN). Mature RNA was isolated and fragmented using the NEBNext Poly(A) mRNA Magnetic Isolation Module (New England Biolabs Inc), following manufacturer’s instructions. The libraries were prepared using the NEBNext Ultra Directional RNA Library Prep Kit (Illumina), following manufacturer’s instructions. The NEBNext Multiplex Oligos from Illumina (Index Primers Set 1, New England Biolabs Inc) was used for the library preparation using the following PCR program: 98°C for 30 sec (1 cycle), 98°C for 10 sec followed by 65°C for 75 sec (8 cycles), and 65°C for 5min. The PCR product (≈50μl) was then purified using 45μl of Oligo dT Beads (Agencourt AMPure XP), followed by three washes with ethanol 80%. After air-drying, the dscDNA was eluted by adding 23μl of 0.1x TE, and the resulting 20μl of the library were collected. Sequencing was run on an Illumina HiScan SQ sequencer, using a high output run and sequencing by synthesis V3 chemistry on a single 100 base pair (bp) run. The sequencing process was monitored in real-time using Sequencing Analysis Viewer v1.8.37 Software, and therefore quality control was performed during and after sequencing. Then, the data was imported into a FASTA file format to perform the analysis.

### 11. RNA Sequencing analysis

#### Data processing

Featurecounts was run to obtain per-gene read counts. We calculated the transcriptome coverage for each sample, defined as the number of unique sequencing reads aligning to annotated transcriptomic regions. Coverage was estimated by multiplying the number of assigned reads (as determined by featureCounts) by the read length and dividing by the estimated total transcriptome length of approximately 76 Mb (from the Ensembl GRCz11_109 GTF file). On average, we obtained a coverage of 12X across all samples, corresponding to an average assignment rate of approximately 65%.”

<u>Differential expression:</u> To identify signatures of ageing, WT brain samples were subjected separately to DESEq2 analysis, comparing the time points 9, 22 and >30 months with the time-point of 2 months. Then, *tert-^/-^* samples at 2, 9 and 22 months were compared with WT at 2 months, to identify telomerase-dependent ageing processes.

<u>Enrichment analysis:</u> Pathway enrichment analysis was performed using the Functional Annotation Bioinformatics Microarray Analysis programme (https://david.ncifcrf.gov/). Pathways identified in both Gene Ontology Biological Process (GO-BP) and KEGG databases were included in the analysis. The top 10 GO-BP and KEGG terms were then plotted into a heatmap using graphpad Prism.

Hypergeometric enrichment analysis:

To assess whether the genes identified through the STEM analysis were over-represented in specific brain cell types, we performed over-representation analysis using Fisher’s Exact Test under the hypergeometric distribution. For each cell type from the previously published adult zebrafish brain single-cell RNA-seq dataset ^65^, we used the authors’ defined gene programmes, consisting of marker genes with adjusted p-values < 0.05. These marker genes were compared to the list of STEM analysis genes to test whether the overlap was greater than expected by chance, relative to the total number of protein-coding genes in the zebrafish genome. A Fisher’s Exact Test was applied for each cell type and resulting p-values were adjusted for multiple comparisons using the Bonferroni correction method.

<u>Mapping to the hallmarks of ageing:</u> We analysed the recent Frenk & Houseley’s paper^67^, which describes the different transcriptional hallmarks of aging. In this paper, the different hallmarks are defined within specific text sections. For each hallmark, we collected groups of keywords that could be within GO term names. The complete list of GO term names was extracted from the file go-basic.obo from the Gene Ontology website (https://geneontology.org/). We then established lists of GO term names for each transcriptional hallmark of aging. The mapping table between the hallmarks, the keywords, and the lists of GO term names is available in the table GO_to_HA-THA_V1.csv.

Zebrafish genes and their associated GO term names were retrieved from the mapping function Gene2GoReader() from the Python library goatools. We built 3 distinct dictionaries for each GO category (Biological Process, Molecular Function, and Cellular Component), where the entrezgene IDs are associated with their GO term names. From the differential analyses, we obtained different lists of gene symbols that must first be mapped to entrezgene ID, using the Python library mygene. Each gene ID is associated with a list of GO term names. To assess a particular transcriptional hallmark of aging to a given gene, we parsed their associated list of GO term names and mapped them to their corresponding hallmark of aging.

Depending on the situation, the attribution of the hallmark of ageing is done as follows: 1) all the GO term names are mapped to the same transcriptional hallmark of ageing, 2) the GO term names are mapped to several transcriptional hallmarks of ageing, the hallmark with the highest number of occurrences is chosen. Otherwise, the gene is assigned with an “ambiguous” hallmark of ageing if 3) none of the GO term names are mapped to any transcriptional hallmarks of ageing, then the gene is not annotated with a transcriptional hallmark of aging.

### 12. cDNA synthesis for Real-time qPCR

cDNA synthesis was performed using the the SuperScript II RT Kit (Thermo Fisher Scientific, USA) kit, following manufacturer’s instructions. Approximately 300ng of each RNA sample was mixed with 0.5 μL of 500 μg/mL random primers, 1 μL 10 mM dNTPs, and sterile, PCR-grade water to a final volume of 12 μL. Samples were heated to 65 °C for 5 minutes, followed by cooling at 4 °C and placement on ice. To each sample, 4 μL First-Strand Buffer, 2 μL 0.1 M DTT, and 1 μL NxGen RNase inhibitor (LGC Biosearch Technologies, Hoddesdon, UK) were added and gently mixed. Samples were heated to 25 °C for 2 minutes, followed by the addition of 1 μL SuperScript™ II RT. Samples were then incubated at 25 °C for 10 minutes, followed by incubation at 42 °C for 50 minutes, before the reaction was inactivated by heating at 70 °C for 15 minutes.

#### Real-time qPCR

cDNA samples from zebrafish brains were analysed by real-time qPCR using SsoAdvanced Universal SYBR Green Supermix (Bio-Rad), forward and reverse primers at a final concentration of 200 nM (Table 1), and cDNA diluted 1:40. Samples were analysed in triplicate. The CFX384 Touch Real-Time PCR Detection System (Bio-Rad) was used to run each real-time qPCR. The program settings were: 95 °C for 30 seconds, followed by 40 cycles of 95 °C for 5 seconds and 58.4 °C for 15 seconds. A melt curve analysis was included in each experiment by increasing the temperature from 65 °C to 95 °C at a 0.5 °C/ 5 second increment. To ensure that the Cq values of the sample cDNA fell within a linear range in which the primer efficiency was between 90-110%, a two-fold serial dilution of pooled cDNA from randomly selected samples were included in each run, with curves containing at least 5 data points. No template (water) and no reverse transcriptase controls were also included in each run.

During analysis, the standard error of each triplicate was calculated and samples with triplicates presenting a standard deviation ≥0.5 were excluded. The fold change expression for each gene was calculated using the ΔΔCt method [1]. Expression of each target gene was normalised to the housekeeper gene (*beta-actin*) and the average fold-change in values for each group compared to the young WT group was calculated.

Graphs and respective statistical analysis were performed using GraphPad Prism 10.4.2. Normality was assessed using the Shapiro-Wilk test and statistical comparisons carried out using a two-way ANOVA, followed by Dunnett’s test for multiple comparisons. A p-value of <0.05 was considered significant.

### 13. Protein extraction and quantification

Fish were culled by Schedule 1 method as previously described. Brain tissue was dissected, transferred to a microcentrifuge tube and immediately snap-frozen in liquid nitrogen. The samples were stored at -80°C until protein extraction.

The tissue was homogenised in 100μl of ultra-pure water using a mechanical homogenizer (VWR International) and RNAse-free disposable pestles (Thermo Fisher Scientific) for approximately 30sec, on ice. Then, the samples were spun down at 4°C, at 2,000rpm, for 5min. After transferring the supernatant to a new tube, samples were stored at -80°C until further use.

Protein extracts were quantified by the bicinchoninic acid (BCA) method, using the Pierce™ BCA Protein Assay Kit (Thermo Fisher Scientific), following manufacturer’s instructions. Briefly, a standard curve was prepared with a range of 0.2-2μm of BSA, along with a blank control (containing ultra-pure water only). A total of 5μl of each protein extract was mixed with 1ml of BCA, in microcentrifugation tubes, and incubated at 37°C for 10min in a water bath. After adding an additional 20μl of copper sulphate, the solution was mixed and incubated at 37°C for 20min. Then, 200μl of each sample (or BSA standard curve) were transferred to a Corning™ Costar™ Brand 96-Well EIA/RIA Plate (Thermo Fisher Scientific), in duplicates, and absorbances were read in a Varioskan Plate Reader at a wavelength of 562nm.

### 14. Chitotriosidase activity assay

Chitotriosidase activity was measured using the protocol previously described by^100^. This method allows assessment of the hydrolysis capacity of 4–methylumbelliferyl–β-D-N, N’, N”- triacetylchitotriose (4-MU-chitotrioside), by chitotriosidase enzyme.

Briefly, a total of 10μl of protein extract per sample was mixed with 100μl of substrate solution and incubated at 28°C for 6 min. A blank control containing 10μl of ultra-pure water was prepared alongside the samples. Substrate solution consisted in 4-MU-chitotrioside (SIGMA-ALDRICH) diluted in McIlvaine solution at pH 5.2 (1.73mg of 4-MU-chitotrioside per 100 ml of McIlvaine solution). To make the McIlvaine solution, stock solutions of 0.1M citric acid (SIGMA-ALDRICH) and 0.2M di-sodium hydrogen phosphate (SIGMA-ALDRICH) were mixed at a ratio of 45% and 55%, respectively, obtaining a final pH of 5.2. After incubation, 1ml of stop solution (0.25M glycine buffer, pH 10.4, containing 0.01% Triton X-100 (SIGMA-ALDRICH) was added to each sample (and blank control), and 200μl of the final solution was transferred to a Corning™ Costar™ Brand 96-Well EIA/RIA Plate (Thermo Fisher Scientific). Duplicates of each sample were used. An extra microcentrifuge tube was prepared with 200μl of standard solution (5μm 1,4-Methylumbelliferone, SIGMA-ALDRICH) and 900μl of stop solution. A volume of 200μl of the final solution was transferred to the 96-well plate, in duplicates. The fluorescence was then read in a Varioskan Plate Reader (excitation: 365nm; emission: 450nm). Finally, the chitotriosidase activity was calculated in nM/h/ml, using Formula 4.

**T**, fluorescence of test samples

**B**, fluorescence of blank

**Std**, standard

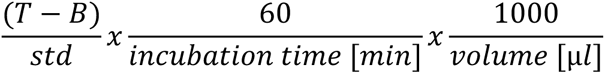

### 15. Blood Brain Barrier (BBB) permeability assay

To test BBB permeability, a total of 5μl of 0.1% IRDye 680RD Dextran (LI-COR Biosciences, Lincoln, NE, USA) diluted in HBSS (Thermo Fisher Scientific) were injected by intraperitoneal injection (IP). Two different sizes of the compound were tested, with the following molecular weights: 4kDa and 70kDa. A group of fish injected with 5μl of HBSS alone was used as a vehicle control group. The fish were culled 3h after IP, and the brains were collected and imaged using a LI-COR Odyssey CLx machine (LI-COR). The brain fluorescence was then quantified using the Image Studio 4.0 Software for Odyssey CLx (LI-COR).

### 16. Mitochondrial complex assays

Homogenised brain tissue was diluted to approximately 0.7mg protein/ml and subjected to 3 cycles of rapid freeze-thawing in liquid nitrogen. All spectrophotometric assays were performed at 28°C in 200µl final volume using a Fluostar Omega plate reader. For assessing complex I activity, the assay buffer was 25mM KH_2_PO_4_, 5mM MgCl_2_, (pH7.2), 3mM KCN, 2.5mg/ml BSA, 50µM ubiquinone, 2µg/ml antimycin A. Complex I activity was determined by measuring the oxidation of nicotinamide adenine dinucleotide reduced disodium salt (NADH) at 340nm with a reference wavelength of 425nm (*ε* = 6.22mM^−1^ cm^−1^). 125µM NADH started the reaction, and 3µg/ml rotenone was used to inhibit the reaction. For assessing Complex II activity, the brain homogenate was incubated for 10min in assay buffer containing 25mM KH_2_PO_4_, 5mM MgCl_2_, (pH 7.2), 3mM KCN, 20 mM succinate. 50µM 2,6-DCPIP, 2µg/ml antimycin A and 3µg/ml rotenone were then added, and baseline measurements recorded. The reaction was started with the addition of 50µM ubiquinone Q1 and complex II activity determined by measuring the reduction of 2,6-dichlorobenzenone-indophenol sodium salt (2,6-DCPIP) at 600nm (*ε* = 19.2mM^−1^ cm^−1^). Complex I and complex II activity rates are normalised to citrate synthase activity.

### 17. Novel tank diving test (NT)

The novel tank diving test (NT) was performed as previously reported by ^93^ and^94^. Briefly, 6 T-75 transparent flasks filled with aquarium water were inserted in the Zantiks LT behavioural unit (Zantiks ltd, Cambridge, UK). After 1h acclimatisation period in the behaviour room, the fish were individually placed into the flasks and allowed to free swim for 5 min. A side camera was used to record the behaviour of the fish for the duration of the experiment. After the test, fish were returned to their home tanks. The fish tracking was used to calculate the time spent and the total distance swam in the top area of the tank (top third of the flask), which were used as measures of anxiety-like behaviour. The more time spent, and the longer distance swam in the top part of the tank indicate lower anxiety levels. Moreover, the total distance swam throughout the entire tank was used as a measure of general motor capacity. This was used as a control to test whether differences in distance swam in the top part of the tank identified between groups was not due to a potential locomotive defect but likely associated with an anxiety-like phenotype.

### 18. Edu Pulse chase experiment and analysis

EdU pulse chase was performed as described ^72, 101^. In brief, fish were fasted for 18 h then anaesthetised in 4% MS-222 (Covetrus, pharmaceutical grade) injected Intraperitoneally (IP) with 5ul 10mM EdU ((Click it EdU Alexa Fluour 647 Imaging Kit (Invitrogen, C10340)) or control solution ((Hanks’ Balanced Salt Solution (HBSS) (Gibco, 14,175–053)) using a 30G 3 ml insulin syringe (BD Micro-Fine U-100 insulin, REF 324,826). Fish were injected once every day for 3 consecutive days at the same time of day (3-day pulse) and culled on day 4 (1 day chase), for paraffin embedding, sectioning and IF (*N* = 4 for each genotype/age).

Immunofluorescent images were processed using ImageJ Fiji (v1.54p) and QuPath (v0.6.0). Maximum intensity Z-projections were generated, with a minimum of three projections analysed per animal. Total cell counts were obtained from Channel 1 (DAPI) using QuPath’s automated cell detection. EdU^+^ cells (Channel 4) were manually counted, and results were expressed as the percentage of EdU^+^ cells per field of view

### 19. Statistical analysis

All the graphs and respective statistical analysis were performed with GraphPad Prism 7.0. Normality was assessed by comparing group variances using either the Brown-Forsythe test or the Shapiro-Wilk normality test. Unpaired t-test or Mann Whitney’s unpaired t-test were used to compare 2 groups, depending on whether the samples presented a Gaussian distribution or not, respectively. For groups with normal distribution, comparisons over-time were performed using ANOVA, followed by Turkey test for multiple comparisons. For groups that failed the normality tests, non-parametric tests were performed, followed by Dunnett’s test for multiple comparisons. A critical value for significance of p<0.05 was used in all the analysis.

## ABBREVIATIONS

AD: Alzheimer’s disease
ALS: amyotrophic lateral sclerosis
ANOVA: analysis of variance
ATH: ageing transcriptional hallmarks
BBB: blood-brain barrier
BCA: bicinchoninic acid
bp: base pair
BSA: bovine serum albumin
Ce: cerebellum
Cent: centromere
CNS: central nervous system
DAPI: 4ʹ,6-diamidino-2-phenylindole
DAVID: database for annotation, visualization, and integrated discovery
DDR: DNA damage response
DEG: differential expression gene
Die: diencephalon
dH2O: distilled water
DNA: deoxyribonucleic acid
Dpf: day post fertilisation
dscDNA: double-stranded cDNA
EDTA: ethylenediaminetetraacetic acid
ENU: N-ethyl-N-nitrosourea
FAM: carboxyfluorescein
FISH: fluorescence *in situ* hybridisation
GEO: gene expression omnibus
GOBP: gene ontology biological pathway
h: hours
HBSS: Hanks buffered solution
IF: immunofluorescence
IP: intraperitoneal injection
IR: irradiation
KCN: potassium cyanide
KEEG: Kyoto encyclopaedia of genes and genomes
KH_2_PO_4_: potassium dihydrogen phosphate
KO: knock-out
LCP1: Lplastin
LMF: Wolfson light microscopy facility
LPS: lipopolysaccharide
MgCl2: magnesium chloride
Min: minutes
MO: medulla oblongata
NaCl: sodium chloride
NADH: nicotinamide adenine dinucleotide reduced disodium salt
NFkβ: nuclear factor kappa β
nm: nanometers
NO: novel object
NPCs: neural progenitor cells
NSCs: neural stem cells
O.C.T.: mounting media
OT: optic tectum
PBS: phosphate buffered saline
PCA: principal component analysis
PCNA: proliferating cell nuclear antigen
PFA: paraformaldehyde
PNA: peptide nucleic acid
RIN: RNA integrity number
RNA: ribonucleic acid
RPM: revolutions per minute
RT: room temperature
SASP: senescence-associated secretory phenotype
sec: seconds
SITraN: Sheffield Institute for Translational Neuroscience
SSC: saline-sodium citrate
TE: Tris-EDTA
Tel: telencephalon
Telo: telomere
Tert: telomerase
TERT: telomerase reverse transcriptase
TNFα: tumour necrosis factor alpha
WT: wild-type
ZIRC: Zebrafish international resource center
4-MU-chitotrioside: 4–methylumbelliferyl–β-D-N, N’, N”-triacetylchitotriose
2,6-DCPIP: 2,6-dichlorobenzenone-indophenol sodium salt

## ACKNOWLEDGEMENTS

We would like to acknowledge and thank Dr Marcus Keatinge for helping with the chitotriosidase activity assay. We would like to acknowledge and thank the University of Sheffield Aquarium staff for the great care they provide to our fish and help with protocols, fish selection and breeding. We would like to acknowledge and thank the Wolfson Light Microscopy Facility staff for their support throughout the years with all our imaging needs and challenges. Finally, we would like to acknowledge John Kemp for critical discussion about transcriptomic data and hypergeometric analysis and statistics.

This work was generously funded by a University of Sheffield PhD studentship to RRM. This research was funded in whole, or in part, by Wellcome [206224/Z/17/Z/WT_/Wellcome Trust/United Kingdom]. For the purpose of open access, the author has applied a CC BY public copyright licence to any Author Accepted Manuscript version arising from this submission. CMH was also funded by a Vice Chancellor’s Fellowship, The University of Sheffield and a VIVENSA grant AISRPG2305\32, which cover PSE and CP salaries.

RS is funded by the BBSRC SwbioDTP and CH by BBSRC (BB/Y002504/1).

MR is funded by the CNRS and the ANR ADAGIO (ANR-20-CE44-0010). Thanks to the Bettencourt Schueller Foundation long-term partnership, this work was partly supported by the CRI Core Research Fellowship to Michael Rera for SB salary.

## CONFLICT OF INTEREST

The authors declare no competing interests.

## AUTHOR CONTRIBUTIONS

*Author contributions according to CRediT:*

**RRM**: conceptualisation, data curation, formal analysis, funding acquisition, investigation, methodology, project administration, validation, visualisation, writing

**SB:** data curation, formal analysis, investigation, methodology, software, validation, visualisation, writing

**PSE:** data curation, formal analysis, investigation, methodology, validation, writing

**RS:** data curation, formal analysis, investigation, methodology, writing

**NH:** data curation, formal analysis, investigation, methodology

**CP:** data curation, formal analysis, investigation, methodology, writing

**GEB:** formal analysis, methodology, software

**OE:** investigation

**NM:** investigation

**MHFW:** investigation

**ZY:** investigation

**SM**: methodology

**AJG:** methodology

**AG:** methodology

**HM:** data curation, formal analysis, funding acquisition, investigation, methodology

**CH:** data curation, formal analysis, funding acquisition, investigation, methodology

**MR:** data curation, formal analysis, funding acquisition, investigation, methodology, project administration, software, validation, visualisation, writing

**CMH:** conceptualisation, data curation, formal analysis, funding acquisition, investigation, methodology, project administration, resources, supervision, validation, writing

## DATA AVAILABILITY STATEMENT

The dataset and source data generated during and/or analysed during the current study is either shared as supplementary and source data or is available from the corresponding author upon reasonable request. The RNA sequencing data from this experiment were deposited in gene expression omnibus (GEO), accession GSE205601.

## Notes

https://www.ncbi.nlm.nih.gov/geo/query/acc.cgi?acc=GSE205601

### Competing Interest Statement

The authors have declared no competing interest.

### Summary of Updates

This is a revised version which we submitted in response to reviewers comments in the journal in which is under revision. It adds new data on proliferation, DNA damage and offers new insights via an additional STEM and hypergeometric enrichment analysis, contextualising our transcriptomic to putative cell types.

