## Supplementary figures and legends for "TELOMERASE DEPLETION ACCELERATES AGEING OF THE ZEBRAFISH BRAIN"

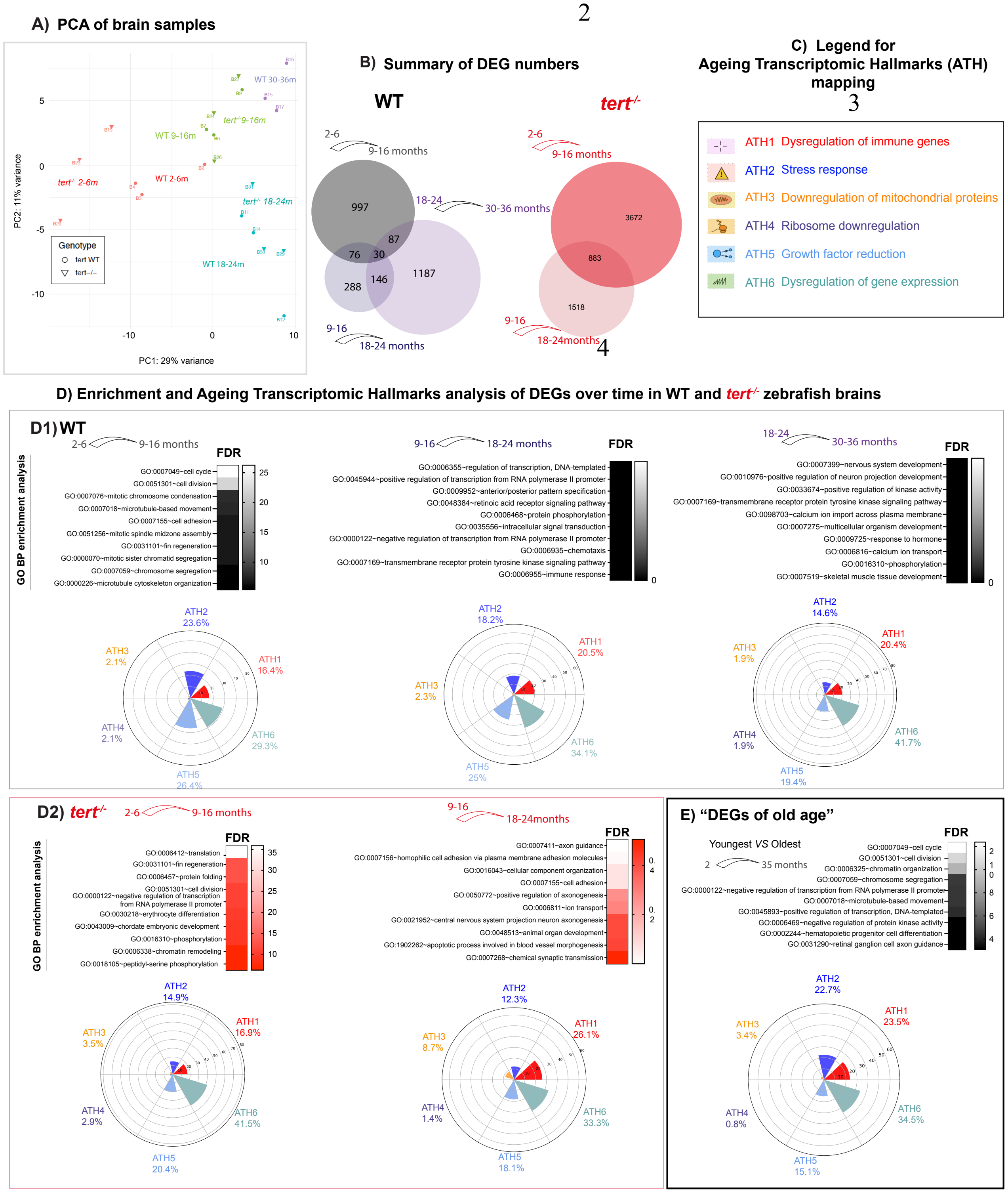


**Supplementary figure 1: Age-associated DEGs are mainly associated with dysregulation of gene expression, dysregulation of immune genes, stress response and growth factor reduction, in both WT and *tert-/-* fish.** (A) Principal component analysis (PCA) plot representing the distribution of gene expression in all the brain samples, by age and genotype, show that the sample cluster mainly by age group. (B) Venn diagrams showing the number of DEGs identified exclusively between each age timepoint, in WT (grey) and *tert-/-* (red) fish. (C) Legend for the ageing transcriptomic hallmarks (ATH) mapping, based on (Frenk & Houseley, 2018). (D) Enrichment analysis and mapping to the ATH of DEGs throughout the lifecourse of WT and *tert*$^{-/-}$ fish. Heatmaps show the top 10 GOBP and KEGG terms identified by enrichment analysis using DAVID software, between every consecutive timepoint, in (D1) WT and (D2) *tert*$^{-/-}$ fish. Plots with the proportion of the DEGs mapped to each ATH show that altered gene expression, stress response, growth factor reduction and immune response are the hallmarks most affected in both (D1) WT and (D2) *tert*$^{-/-}$; and alteration in mitochondrial proteins is affected in old *tert*$^{-/-}$ but not WT fish. (E) Enrichment analysis and mapping to ATH in DEGs of "old age" (2 vs >30 months in WT fish) show similar results than those identified between specific age groups.

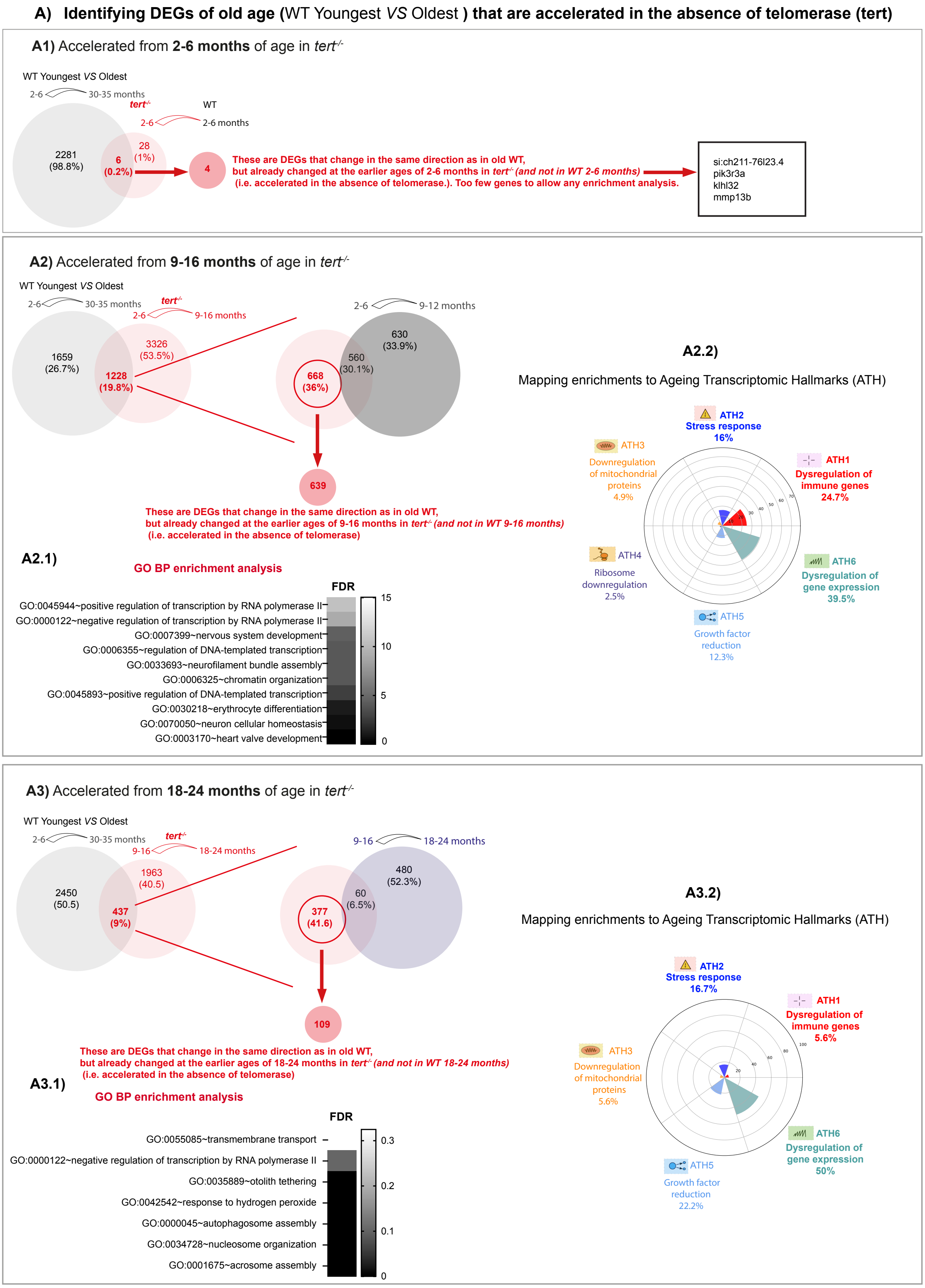


**Supplementary figure 2: DEGs of "old age" that altered by telomerase depletion, and respective enrichment analysis and mapping to ATH.** (A) Venn diagrams show the number of DEGs between every consecutive timepoint in

WT and highlight how many DEGs, within those, are already altered in *tert*$^{-/-}$ at an earlier timepoint, in particular at (A1) 2-6 (young), (A2) 9-16 (young adult) and (A3) 18-24 (old adult) months of age. Enrichment analysis and ATH mapping of such DEGs are shown in heatmaps and proportion plots, respectively, for (A2.1, A2.2) young and (A3.1, A3.2) old adult *tert*$^{-/-}$ fish.

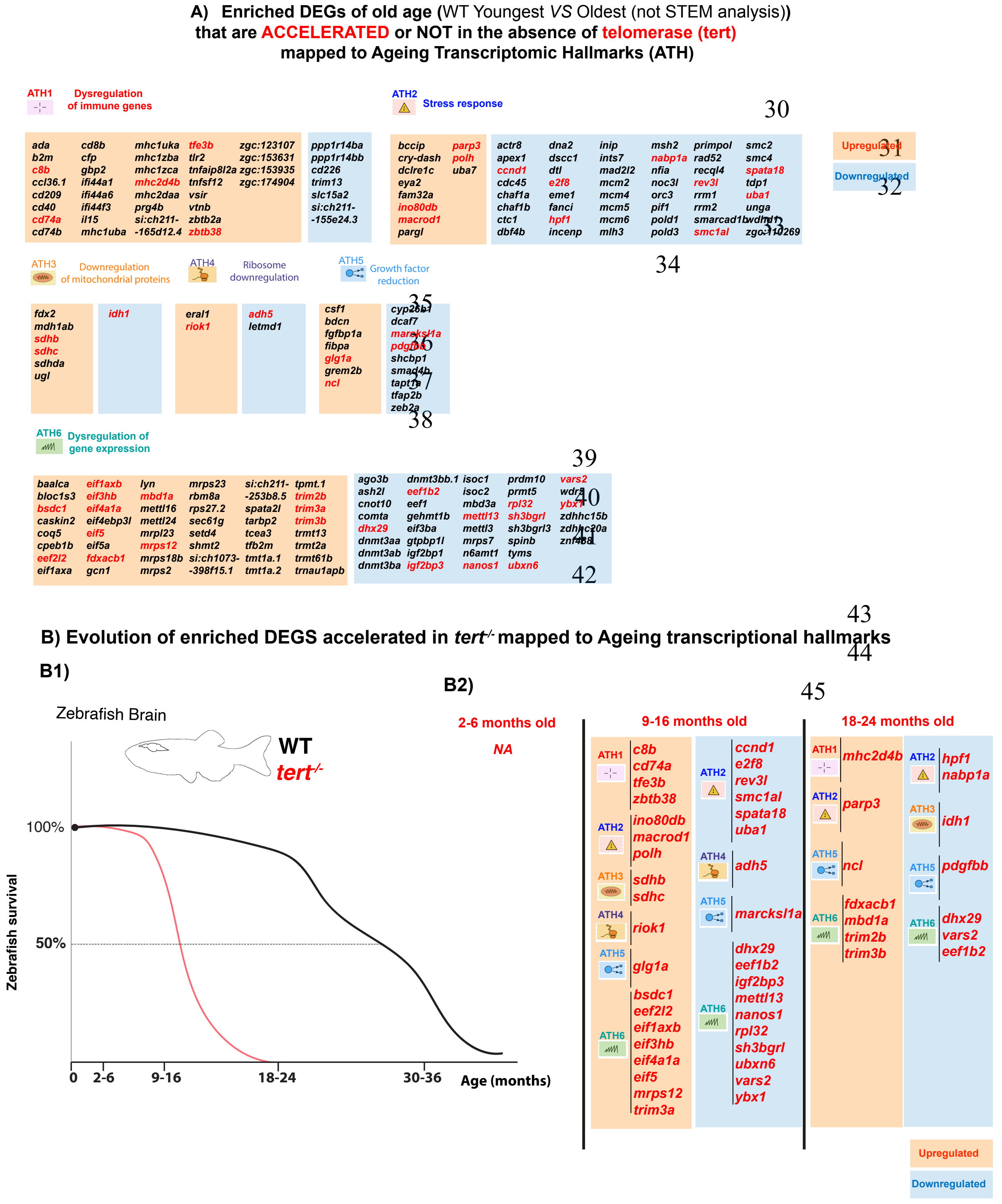


**Supplementary figure 3: List of enriched DEGs of "old age" that are accelerated or not in the absence of telomerase (*tert*$^{-/-}$ fish) and that were mapped to ATH.** (A) Tables show the lists of DEGs mapped to each ATH; up-regulated genes in orange and down-regulated genes in blue. The genes that are altered with natural ageing (WT young vs old) and that are accelerated in the absence of telomerase (*tert*$^{-/-}$ fish) are highlighted in red, whereas the genes altered with natural ageing but that are not accelerated in *tert*$^{-/-}$ fish are in black. (B) Evolution of enriched DEGs accelerated in *tert*$^{-/-}$ fish and that were mapped to ATH. (B1) Survival plot showing the early and late stages of ageing in WT fish compared to *tert*$^{-/-}$ fish. .(B2) List of DEGs of "old age" that are altered at a younger age in *tert*$^{-/-}$ fish and that were mapped to ATH, per age group Up-regulated genes in orange and down-regulated genes in blue.

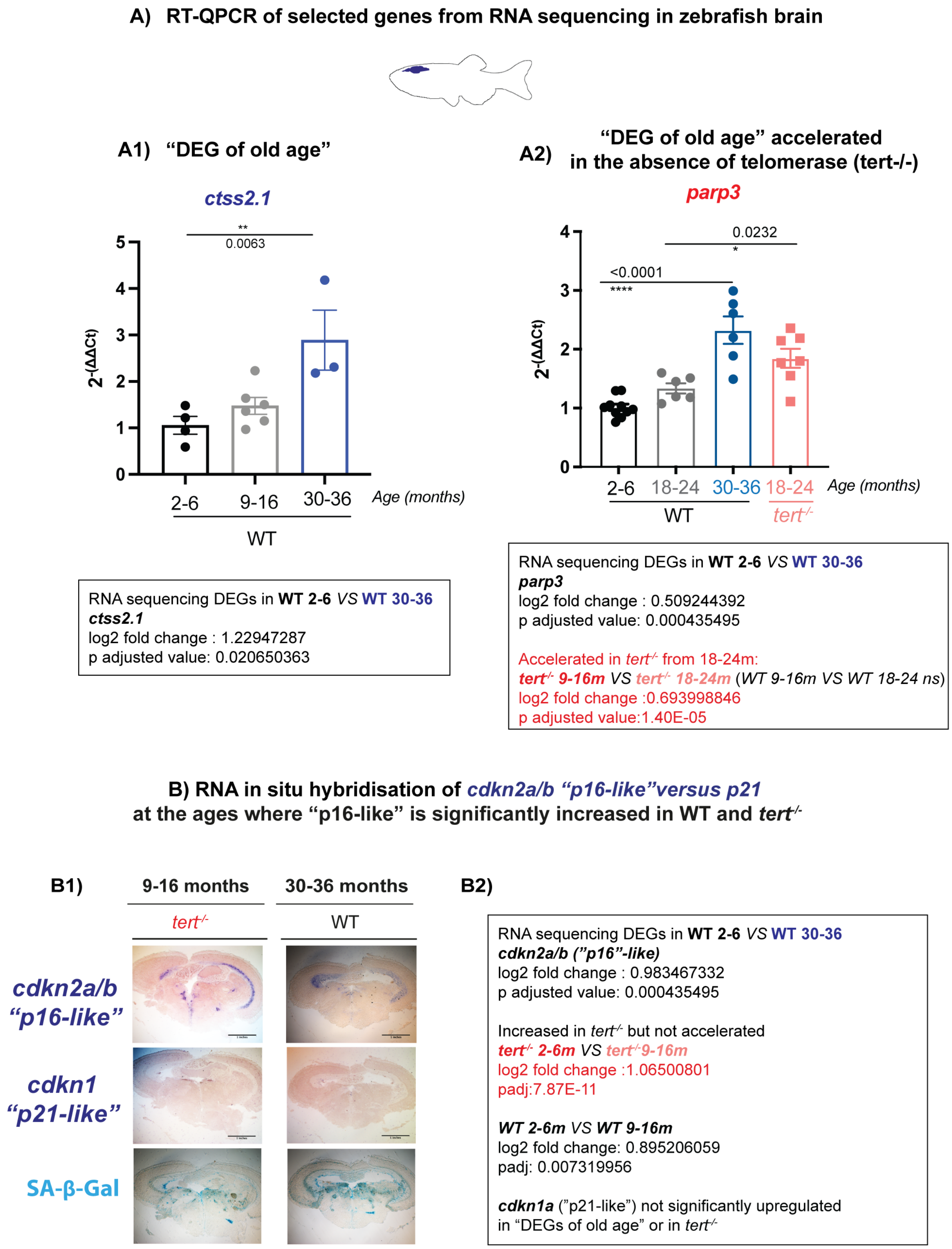


**Supplementary figure 4: Further validation of RNA sequencing data. Further validation of RNA sequencing data.** (A) RT-QPCR of selected genes from RNA sequencing in whole zebrafish brain: (A1) *ctss2.1*, increased with natural ageing;

and (A2) *parp3*, increased with natural ageing and accelerated in *tert*$^{-/-}$ fish. (B1) Representative images of RNA *in situ* hybridisation of *cdkn2a/b* ("*p16-like*") and *cdkn1* ("*p21-like*") in adjacent brain sections, in the same brain regions were SA-β-Gal is observed in old WT and *tert*$^{-/-}$ fish.(B2) Expression of *cdkn2a/b* and *cdkn1a* genes detected by RNA sequencing in WT and *tert*$^{-/-}$ fish. N=3-9. Each dot represents one animal. Bar errors represent the SEM. * <0.05; ** <0.01; *** <0.001.

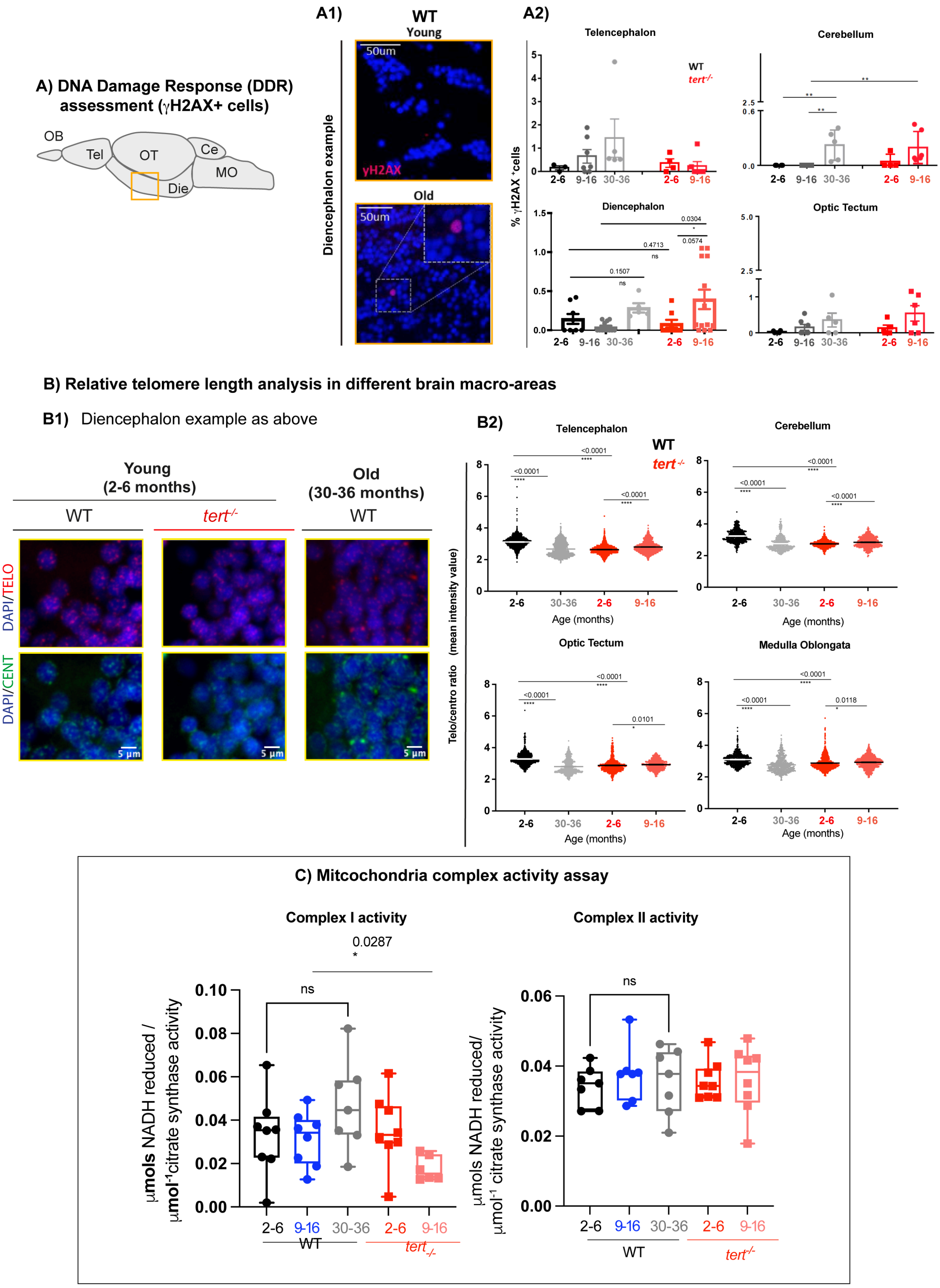

**Supplementary figure 5: Telomerase (tert) depletion accelerates ageing-associated stress response in the zebrafish brain, per macro areas.** (A) DNA damage response (DDR) assessment in brain macro areas, in longitudinal paraffin sections (yellow square highlights the region from where the diencephalon example image was acquired): (A1) representative image from γH2AX staining in the brain (diencephalon used as an example), and (A2) respective quantifications in telencephalon, cerebellum, diencephalon and optic tectum. (B) Relative telomere length analysis in brain macro areas, in coronal paraffin sections: (B1) Representative images from telomere and centromere staining (by FISH) in young and old WT, and young $tert^{-/-}$ brain tissue (diencephalon as an example) and (B2) quantifications of telomere/centromere ratio in the telencephalon, cerebellum, optic tectum, and medulla oblongata.. (C) Assessment of mitochondria complex I and complex II activity in whole brain of WT and $tert^{-/-}$ fish at different timepoints. N=4-9. (A2, C) Each dot represents one animal. (B2) Each dot represents a cell. Bar errors represent the SEM. * <0.05; ** <0.01; *** <0.001.

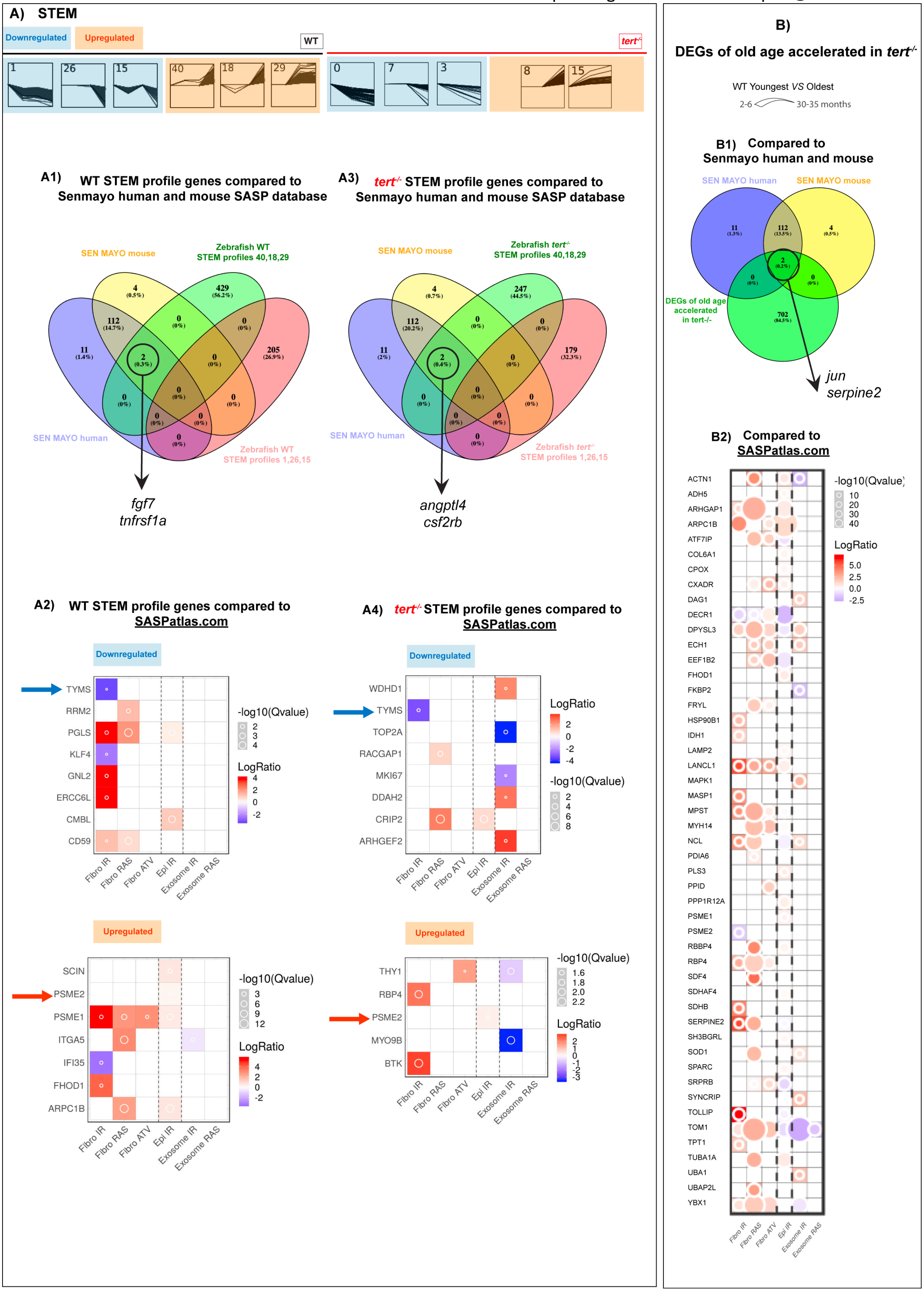


82 **Supplementary figure 6: SASP candidates and telomerase dependencies in aged zebrafish brains. A1**) List of
83 putative SASP molecules identified in the RNA sequencing data of significantly up-regulated DEGs in WT young (c.2

months) *versus* old (>30 months), when searched using the saspatlas.com database. Red arrows point to telomerase Tert-dependent putative SASP molecules, i.e., up-regulated at earlier ages in *tert-/-* zebrafish as identified in A2). Black rectangles highlight that 8/13 of these putative SASP factors are also upregulated in RAS-induced senescence of fibroblasts in vitro, which is the type of senescence with more SASP factors in common with aged zebrafish brain, out of the ones available in saspatlas.com. **A2**) Ven diagram comparing the putative SASP factors of WT aged zebrafish brain with the previously identified tert-dependent DEGs (see Fig 1 and supplementary data source files)

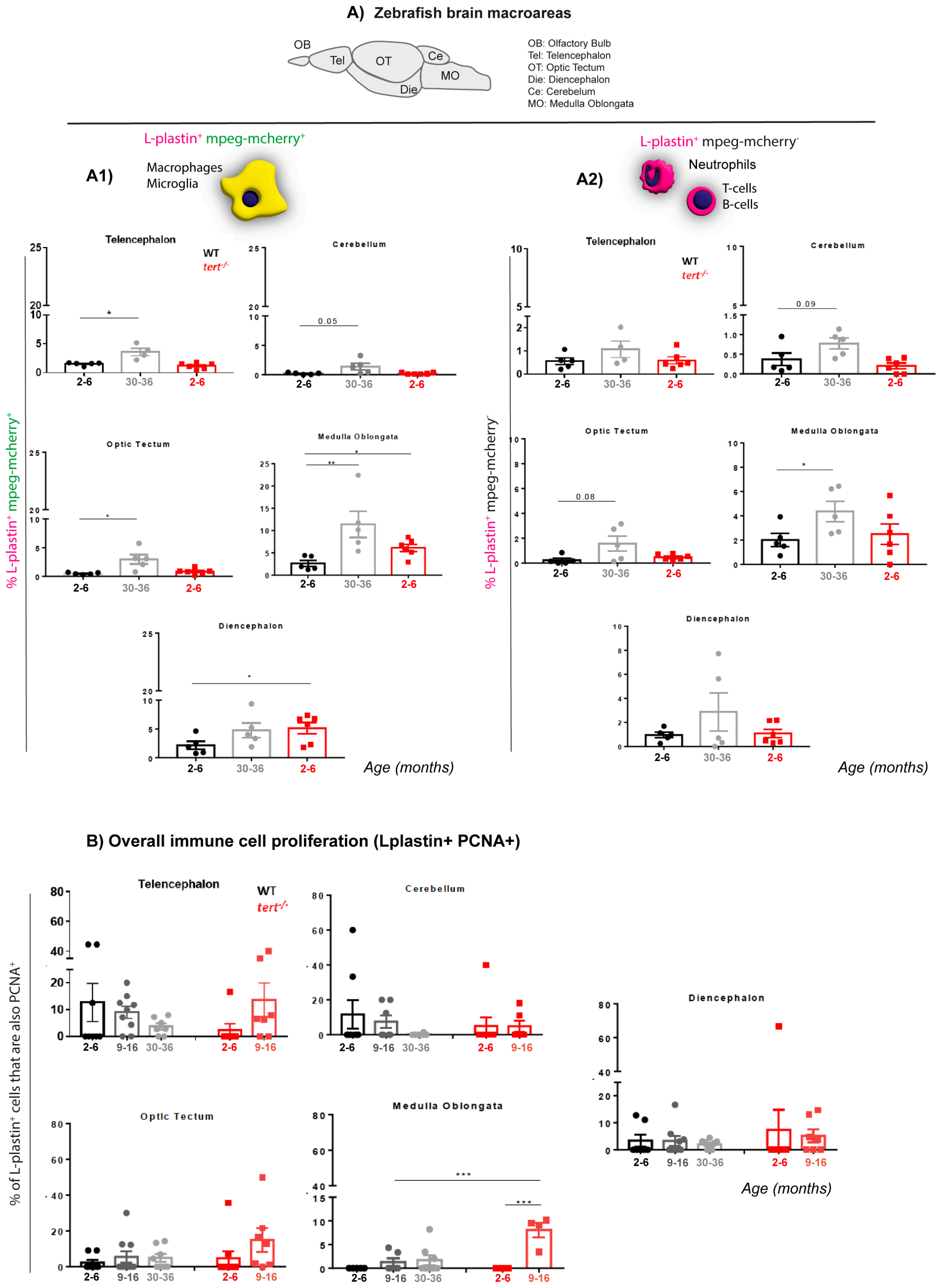

**Supplementary figure 7: Distribution of immune cells in the zebrafish brain with ageing, per macro areas.** (A1) Quantification of macrophages and microglia cells (Lplastin$^+$*; mpeg*-mCherry$^+$ cells) and (A2) quantification of putative T and B cells (Lplastin$^+$*; mpeg*-mCherry$^-$ cells) in the telencephalon, cerebellum, optic tectum, medulla oblongata and diencephalon of WT and *tert*$^{-/-}$ fish. (B) Quantifications of immune cell proliferation (Lplastin$^+$; PCNA$^+$ cells) in the different macro areas of the brain (telencephalon, cerebellum, optic tectum, medulla oblongata and diencephalon), in WT and *tert*$^{-/-}$ fish. N=4-9. Each dot represents one animal. Bar errors represent the SEM. * <0.05; ** <0.01; *** <0.001.
