## Supplementary figures and images for "TELOMERASE DEPLETION ACCELERATES AGEING OF THE ZEBRAFISH BRAIN"

### brain_MT_profiles.png

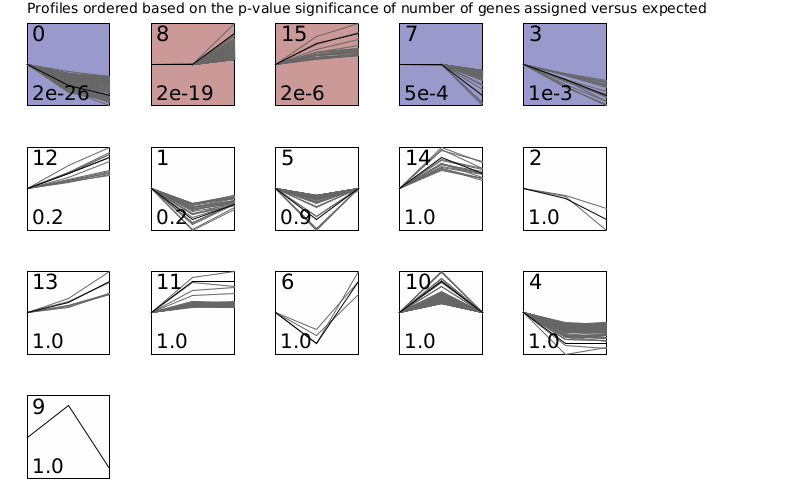

### brain_WT_profiles.png

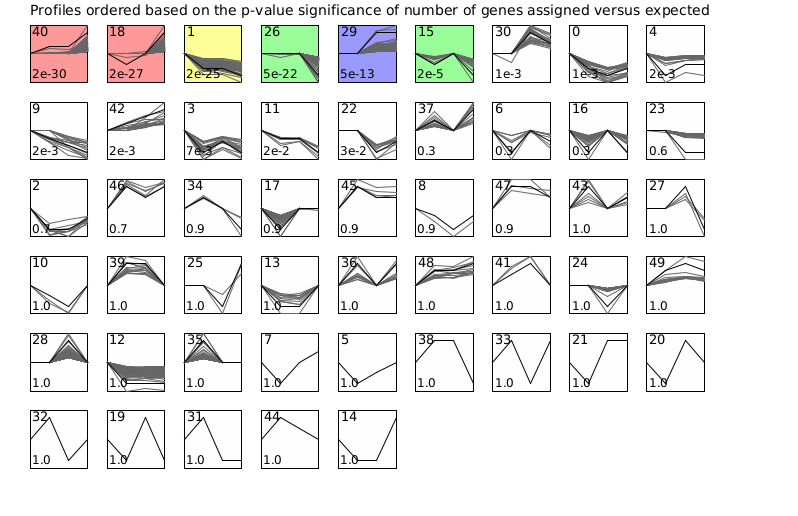
